# Multi-context genetic modeling of transcriptional regulation resolves novel disease loci

**DOI:** 10.1101/2021.09.23.461579

**Authors:** Mike Thompson, Mary Grace Gordon, Andrew Lu, Anchit Tandon, Eran Halperin, Alexander Gusev, Chun Jimmie Ye, Brunilda Balliu, Noah Zaitlen

## Abstract

A majority of the variants identified in genome-wide association studies fall in non-coding regions of the genome, indicating their mechanism of impact is mediated via gene expression. Leveraging this hypothesis, transcriptome-wide association studies (TWAS) have assisted in both the interpretation and discovery of additional genes associated with complex traits. However, existing methods for conducting TWAS do not take full advantage of the intra-individual correlation inherently present in multi-context expression studies and do not properly adjust for multiple testing across contexts. We developed CONTENT— a computationally efficient method with proper cross-context false discovery correction that leverages correlation structure across contexts to improve power and generate context-specific and context-shared components of expression. We applied CONTENT to bulk multi-tissue and single-cell RNA-seq data sets and show that CONTENT leads to a 42% (bulk) and 110% (single cell) increase in the number of genetically predicted genes relative to previous approaches. Interestingly, we find the context-specific component of expression comprises 30% of heritability in tissue-level bulk data and 75% in single-cell data, consistent with cell type heterogeneity in bulk tissue. In the context of TWAS, CONTENT increased the number of gene-phenotype associations discovered by over 47% relative to previous methods across 22 complex traits.

## 1 Introduction

A large portion of the signal discovered in genome-wide associations studies (GWAS) has been localized to non-coding regions [1]. In light of this, researchers have developed post-GWAS approaches to elucidate the functional consequences of variants and their impact on the etiology of traits [2]. One notable approach has been to generate genetic predictors of gene expression and leverage these predictors with GWAS data to associate genes with traits of interest[3, 4]. These transcriptome-wide association studies (TWAS) have not only shown great promise in terms of discovery and interpretation of association signals but have also helped prioritize potentially causal genes for complex diseases [5]. Nonetheless, methods like TWAS are limited by the accuracy and power of the genetic predictors generated in training datasets [6–11].

The original TWAS methodology builds genetic predictors of expression on a context-by-context basis. For example, in a study with RNA-seq and genotypes collected across multiple tissues, the expression of each tissue would be modeled independently [3, 4]. More recent methods model multiple contexts simultaneously and leverage the sharing of genetic effects across contexts [8–10, 12]. However, these approaches do not maximize predictive power because they ignore the intra-individual correlation of gene expression across contexts inherent to studies with repeated sampling, e.g., the Genotype-Tissue Expression (GTEx) project [13] (Figure S1) or single-cell RNA-Sequencing (scRNA-Seq) experiments (Figure S2). Moreover, they build predictors which are mixtures of both context-specific and context-shared (pleiotropic) genetic effects, making it difficult to distinguish the relevant contexts for a disease gene, and are often computationally inefficient [9]. A recent approach by Wheeler et al. [14] does model correlated intra-individual noise with a linear-mixed model, but does not produce combined predictions of expression, reducing overall power. Finally, existing methods employ multiple testing strategies that either fail to control for all tests performed, (e.g., by controlling the false discovery rate (FDR) within each context separately [4, 15]), or act too stringently (e.g., by using Bonferroni adjustment across all contexts [15]). Together, these shortcomings reduce power and interpretability of TWAS.

Here, we introduce CONTENT—CONtexT spEcific geNeTics— a novel method that leverages the correlation structure of multi-context studies to efficiently and powerfully generate genetic predictors of gene expression. Briefly, CONTENT decomposes the gene expression of each individual across contexts into context-shared and context-specific components [16], builds genetic predictors for each component separately, and creates a final predictor using both components. To identify genes with significant disease associations, CONTENT employs a hierarchical testing procedure (termed “hFDR”; see Figure S3) [17, 18]. CONTENT has several advantages over existing methods. First, it explicitly accounts for intraindividual correlation across contexts, boosting prediction performance. Second, by building specific and shared predictors, it can distinguish context-shared from context-specific genetic components of gene expression and disease. Third, it employs a recently developed hierarchical testing procedure [18] to not only adequately control the FDR across and within contexts, but boost power in cases where a gene has a significant association to disease in multiple contexts. Fourth, this adjustment procedure allows for inclusion of other TWAS predictors [3, 4, 8–10, 12], enabling approaches to be complementary in discovering associations. Finally, CONTENT is orders of magnitude more computationally efficient than several previous approaches.

We evaluated the performance of CONTENT over simulated data sets, GTEx[2, 11, 13], and a single-cell RNA-Seq data set[19, 20]. We show in simulations that CONTENT captures a greater proportion of the heritable component of expression than previous methods (at minimum over 22% more), and that CONTENT successfully distinguishes the specific and shared components of genetic variability on expression. In applications to GTEx, CONTENT improved over previous context-by-context methods both in the number of genes with a significant heritable component (average 42% increase in significant gene-tissue pairs discovered) as well as the proportion of variability explained by the heritable component (average increase of 28%) [3, 4]. Consistent with complex cell type heterogeneity within tissues [21–24], we find that in applications to the single-cell data, genetic predictors at the cell type level have substantially more context-specific heritability than the tissue-level models. We then performed TWAS across 22 phenotypes using weights trained on GTEx and scRNA and found that CONTENT discovered over 47% additional significantly associated genes. We provide CONTENT gene expression weights for both GTEx and the single-cell dataset at TWAShub (http://twas-hub.org/).

## 2 Results

### Methods overview

We developed CONTENT, a method for generating genetic predictors of gene expression across contexts for use in downstream applications such as TWAS. Briefly, for each individual, CONTENT leverages our recently developed FastGxC method [16] to decompose the gene expression across *C* contexts into one context-shared component and *C* context-specific components. Next, CONTENT builds genetic predictors for the shared component and each of the *C* context-specific components of expression using penalized regression. We refer to these predictors as the CONTENT(Shared) and CONTENT(Specific) models. In addition, CONTENT generates genetic predictors of the total expression in each context by combining the context-shared and context-specific genetic predictors with linear regression. We refer to these predictors as the CONTENT(Full) models. A given gene may have CONTENT(Specific), CONTENT(Shared), and/or CONTENT(Full) models depending on the architecture of genetic effects.

We residualized the expression of each gene in each context over their corresponding covariates (e.g. PEER factors, age, sex, batch information) prior to decomposing and then fitting an elastic net with double ten-fold cross-validation for both CONTENT(Shared) and CONTENT(Specific). We examined the number of significantly predicted genes as well as the prediction accuracy (in terms of adjusted *R*^2^) between the cross-validation-predicted and true gene expression per gene-context pair. To properly control the FDR for each method across contexts and genes, we employed a hierarchical FDR correction [17, 18] (Figure S3 and Methods). We note that groups of contexts may comprise additional sources of pleiotropy (e.g. in GTEx the group of brain tissues may have their own shared effects in addition to the overall tissue-shared effects). The decomposition of CONTENT is flexible and can account for both levels of pleiotropy among contexts (see Supplementary Methods).

### CONTENT is powerful and well-calibrated in simulated data

We evaluate the prediction accuracy of CONTENT in a series of simulations and compare its performance to the context-by-context approach[3, 4], which builds predictors by fitting an elastic net in each context separately, as well as UTMOST[9], which builds predictors over all contexts simultaneously using a group LASSO penalty. Implicitly, we compare to the method from [14] which decomposes expression into orthogonal context-shared and context-specific components, as the CONTENT(Shared) and CONTENT(Specific) models are related to these components (See Methods). We omit comparison to other TWAS methods as many of them are built on the same framework as the context-by-context approach, or require external data, such as curated DNase I hypersensitivity measurements [8, 10, 12].

We used simulation parameters from GTEx, the largest multi-context eQTL study to-date, as a guideline. Specifically, we generated gene expression and genotype data such that context-specific genetic effects mostly lie on the same loci as context-shared eQTLs, and context-specific eQTLs without context-shared effects are rare [2, 16]. Intuitively, this framework assumes that, most often, SNPs affect expression of a gene in all contexts, but to a different extent in each context (rather than, for example, acting as an eQTL in only a single context). We varied the proportion of contexts with context-specific heritability, the number of context-specific eQTLs without a context-shared effect, the number of causal SNPs, and the intra-individual residual correlation while keeping the number of genes (1000), contexts (20), *cis*-SNPs (500) and the proportion of context-shared and context-specific heritability constant (.3 and .1 respectively).

Throughout our simulations, CONTENT significantly outperformed the context-by-context and UTMOST approaches in terms of prediction accuracy of the total genetic contribution to expression variability (Figures 2A, S4). The average increase in adjusted *R*^2^ between the true genetic component of expression and the CONTENT(Full) predictor was .22 over UTMOST (p<2e-16 paired two-way t-test) and .48 over the context-by-context approach (p<2e-16 paired two-way t-test). Across nearly the entirety of parameter settings, CONTENT generated context-specific components that were uncorrelated with the true context-shared components (mean adjusted *R*^2^=.023, and vice versa .026; Figure 2B,C). This property is central to the objective as it reduces confounding from pleiotropy in downstream applications such as context fine-mapping. As expected, the previous methods failed to disentangle the context-specific and context-shared components (Figure 2B,C), since they were not developed with this property in mind. Our results were consistent under different values of the simulation parameters (Figures S5, S6, S7, S8).

**Figure 1.**
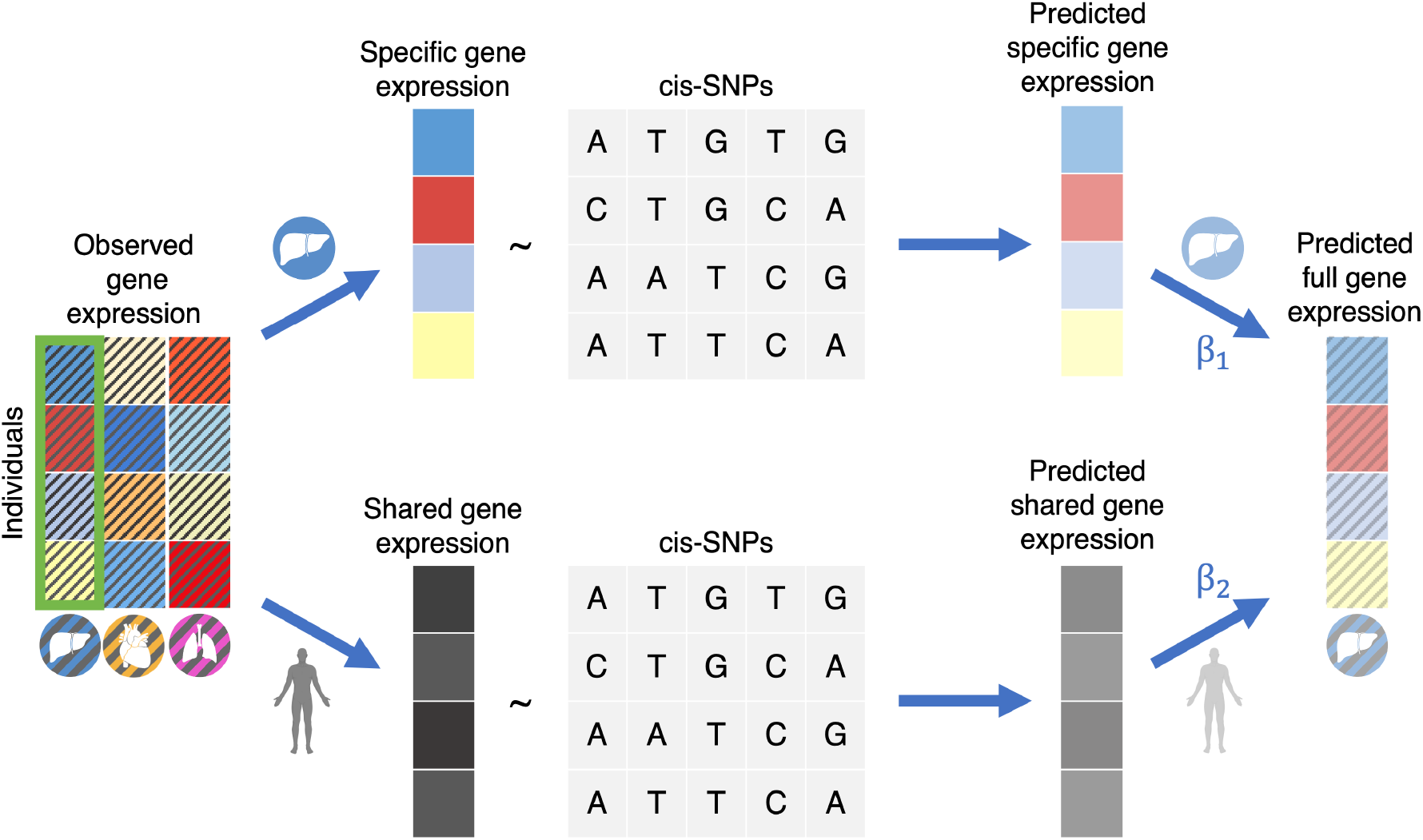
An overview of the CONTENT approach. CONTENT first decomposes the observed expression for each individual into context-specific and context-shared components following [16]. Then, CONTENT fits predictors for the context-shared component of expression as well as each context-specific component of expression (e.g., liver). Finally, for a given context, CONTENT combines the genetically predicted components into the full model using a simple regression.

**Figure 2.**
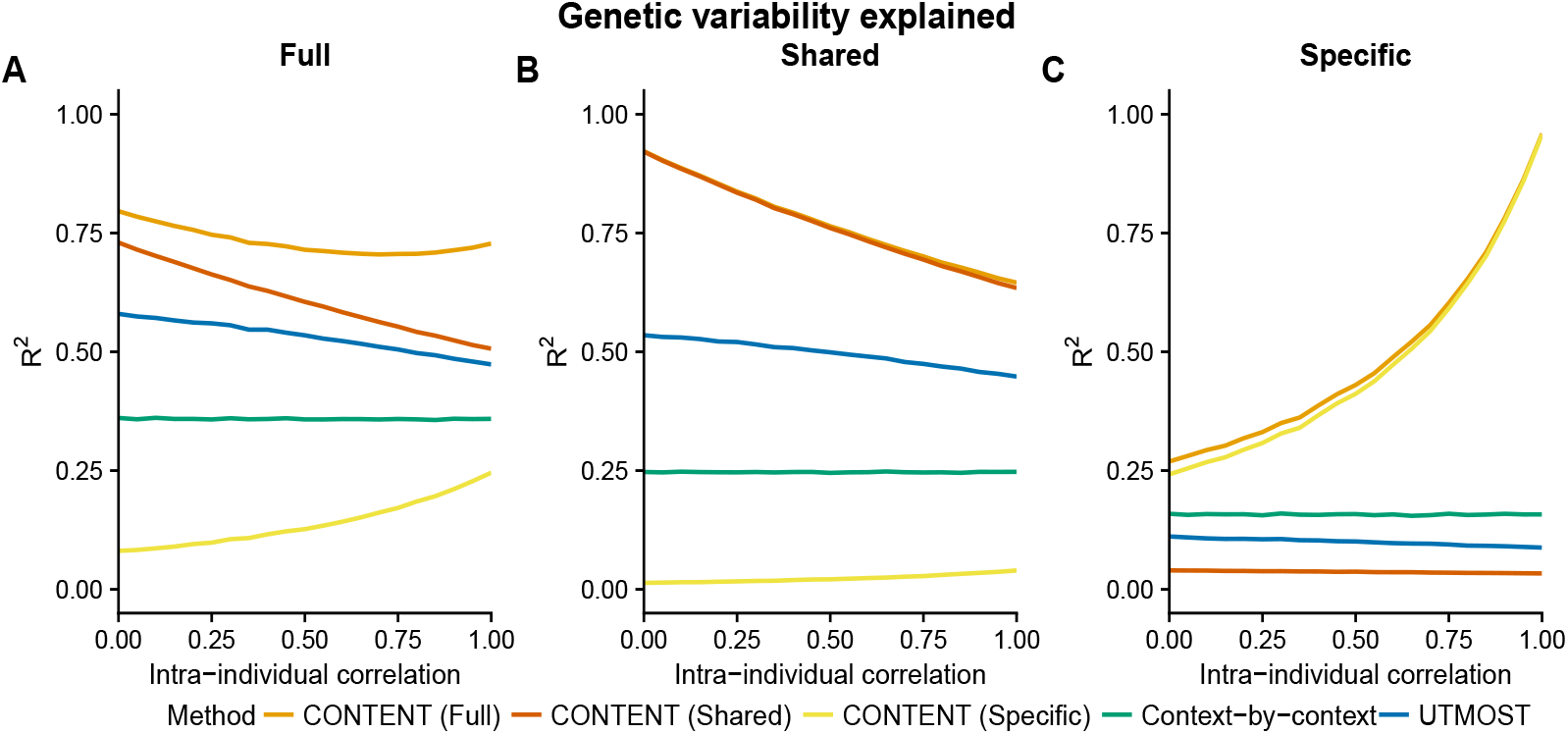
CONTENT is powerful and well-calibrated in simulated data. Accuracy of each method to predict the genetically regulated gene expression of each gene-context pair for different correlations of intra-individual noise across contexts. Mean adjusted *R*^2^ across contexts between the true (A) full (context-specific + context-shared), (B) shared, and (C) specific genetic components of expression and the predicted component for each method and for different levels of intra individual correlation. The context-by-context approach and UTMOST output only a single predictor, and we show the variability captured by this predictor for each component of expression. CONTENT, however, generates predictors for all three components of expression, and notably, CONTENT(Specific) and CONTENT(Shared) capture their intended component of expression without capturing the opposite (i.e. the predictor for CONTENT(Specific) is uncorrelated with the true shared component of expression and vice versa). We show here the accuracy for each component and method on gene-contexts with both context-shared and context-specific effects, but show in Figure S4 the accuracy for all gene-contexts pairs.

### CONTENT improves prediction accuracy over previous methods in the GTEx and CLUES datasets

We next evaluated CONTENT, the context-by-context approach, and UTMOST in terms of prediction accuracy and power across 22,447 genes measured in 48 tissues of 519 European individuals in the bulk RNA-seq GTEx data set [2, 11, 13]. Due to computational issues (Figure S9), UTMOST was examined only on 22,307 genes rather than the entire data set of 22,447 genes. We show a comparison on this smaller set of genes in Figure S10. We also examined, for the first time in a large-scale TWAS context, a single-cell RNAseq data set from the California Lupus Epidemiology Study (CLUES) [19, 20]. The CLUES data set contained 9,592 genes measured in 9 cell types in peripheral blood from 90 individuals.

In GTEx, CONTENT identified more gene-tissue pairs with a significantly predictable genetic component of expression (278,101 over 20,506 genes) than the context-by-context approach (195,607 over 17,723 genes) and UTMOST (167,865 over 11,442 genes) at an hFDR of 5%. We also compared the performance of each method on the union of genes that were significantly predicted (hFDR ≤ 5%) by at least one method. As CONTENT can generate up to three models (Shared, Specific, Full) for a given gene-tissue pair, and because each gene may have its own unique architecture (i.e. different proportions of specific or shared heritability), we selected the model that achieved the greatest cross-validated adjusted *R*^2^. CONTENT greatly outperformed the context-by-context and UTMOST approaches across all tissues (average 28% and 22% increase in adjusted *R*^2^ across tissues and genes; Figures 3, S10). Further, for genes with significant CONTENT(Shared), CONTENT(Specific), and CONTENT(Full) predictors, prediction accuracy increases substantially with the addition of the context-specific component to the context-shared component (average gain of CONTENT(Full) over CONTENT(Shared) adj. *R*^2^ of 55.92%), emphasizing the need to extend previous approaches[14] with CONTENT(Full) to build a powerful predictor.

**Figure 3.**
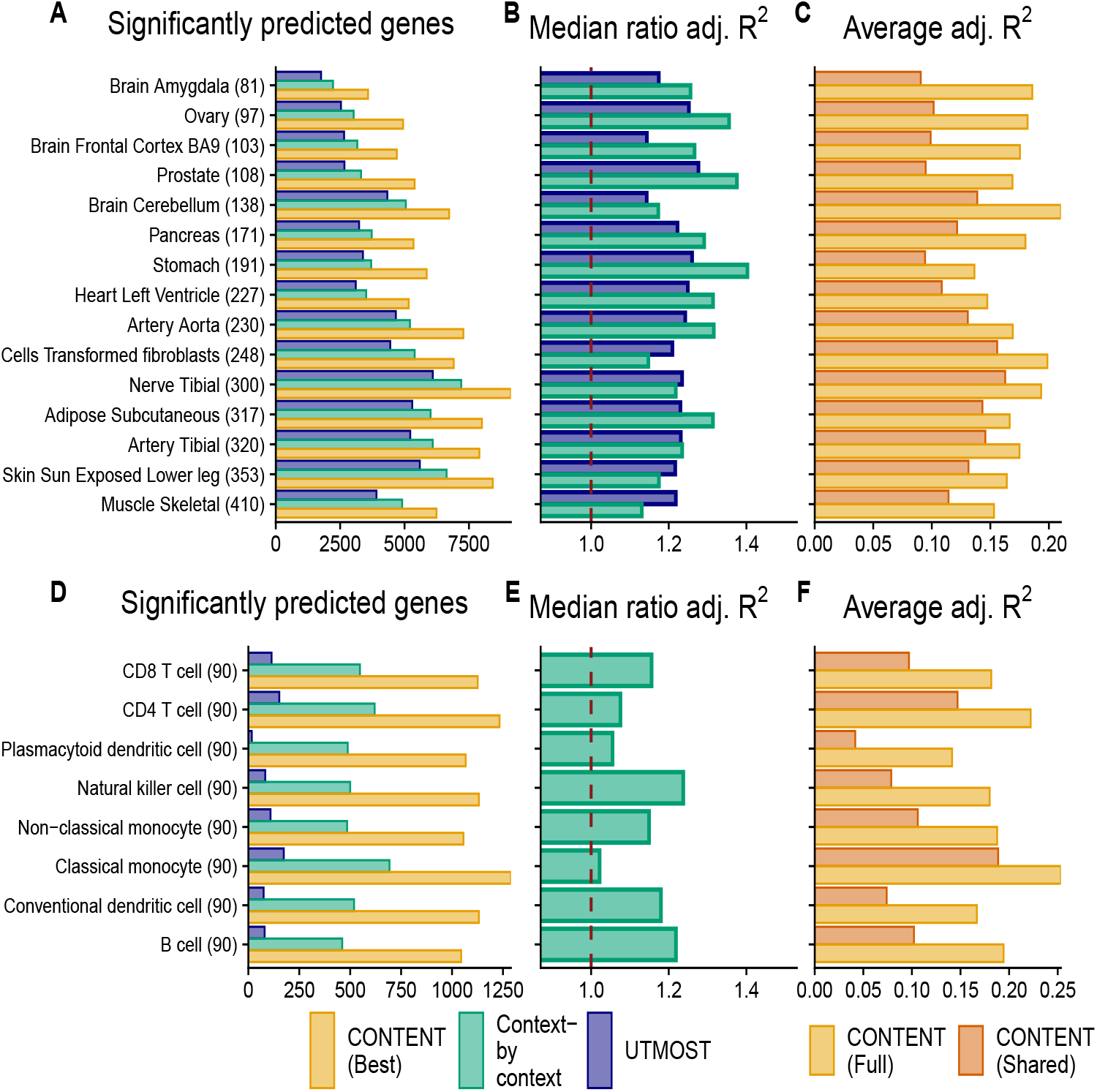
CONTENT outperforms existing approaches in the GTEx and scRNA-seq CLUES datasets. (A,D) Number of genes with a significantly predictable component (hFDR ≤ 5%) in GTEx (A) and CLUES (D); the sample sizes for each context are included in parentheses. (B,E) Ratio of expression prediction accuracy (adjusted *R*^2^) of the best-performing cross-validated CONTENT model over the context-by-context (green) and UTMOST (blue) approaches (median across all genes significantly predicted by at least either method). Numbers above one indicate higher adjusted *R*^2^ and thus prediction accuracy for CONTENT. (C,F) Prediction accuracy of CONTENT(Full) and CONTENT(Shared) when a gene-tissue has a significant shared, specific, and full model.

Within the single-cell CLUES data set, CONTENT again outperformed the context-by-context (in this case, cell type-by-cell type) and UTMOST approaches, discovering 9,080 heritable gene-cell type pairs (5,067 genes) whereas the context-by-context model and UTMOST found 4,314 (2,355 genes) and 804 (288 genes) respectively. The average improvement in adjusted *R*^2^ of CONTENT over the context-by-context model was 13.6%. In gene-cell type pairs with significant CONTENT(Full), CONTENT(Specific), and CONTENT(Shared) models, CONTENT(Full) improved the adjusted *R*^2^ over CONTENT(Shared) by 104.09%. Once more, the improvement in variability explained when including both the cell type-specific and cell type-shared components highlights the need to consider both components simultaneously when building a predictor.

### CONTENT discovers significant context-specific components of expression in bulk multi--tissue and single-cell datasets

Given the ability of CONTENT to disentangle context-shared and context-specific variability, we examined the context-specific components of expression in GTEx and CLUES. In GTEx, CONTENT discovered 128,985 gene-tissue pairs (19,765 genes) with a significant context-specific genetic component of expression (Figures 4, S11). As with previous reports [16, 25], we found that testis was the tissue with the greatest number of tissue-specific genetic components. Nonetheless, we observe that the tissues with larger sample sizes more frequently had significant context-specific components. Consistent with previous works that have discovered extensive eQTL sharing across tissues [2, 25, 26], we found that in gene-tissue pairs with a CONTENT(Full) model, the variability explained was dominated by CONTENT(Shared) model—across tissues, the context-shared component explained on average 70% of the variability explained by CONTENT(Full).

**Figure 4.**
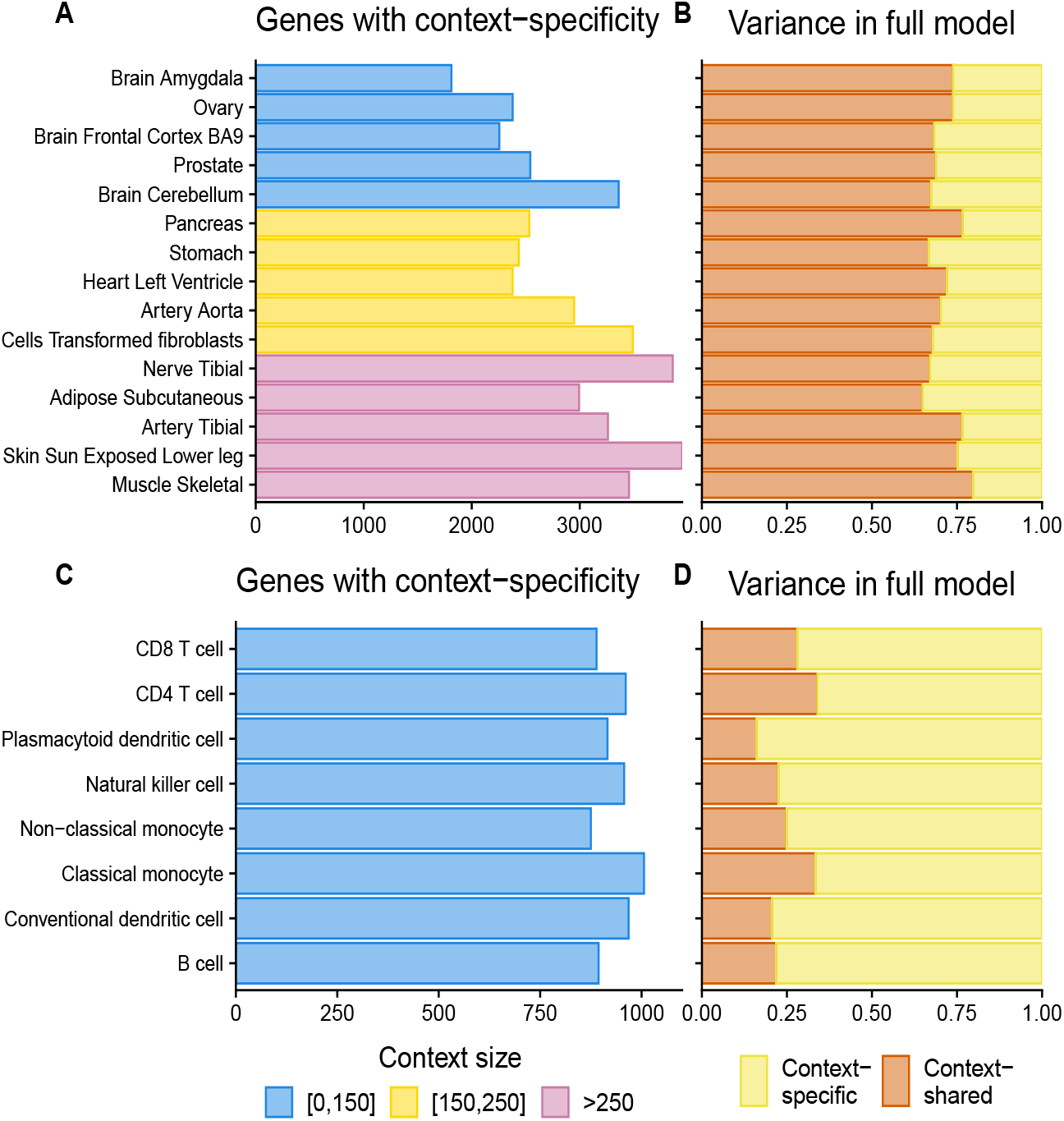
Contribution of context-specific genetic regulation in GTEx and CLUES. (A,C) Number of genes with a significant (FDR 5%) CONTENT(Specific) model of expression in GTEx (A) and CLUES (C). Color indicates sample size of context. (B,D) Proportion of expression variance of CONTENT(Full) explained by CONTENT(Specific) and CONTENT(Shared) for genes with a significant CONTENT(Full) model.

In the CLUES data set, CONTENT discovered 7,466 gene-cell type pairs (4,658 genes) with a significant cell type-specific component of expression (hFDR ≤ 5%). We found that all cell types had a similar number of cell type-specific components, and emphasize that the sample size across all cell types was equivalent. Interestingly, in genes with a CONTENT(Full) model, the variability was often dominated by the cell type-specific variability (average 75% of the explained variability), unlike GTEx, in which the average tissue-specific variability explained only 30% of total variance. Consequently, we found that within the 20,433 genes in GTEx with any genetic component, 51.50% (10,522) had a significant shared component, whereas of the 5,067 genes in CLUES with a genetic component, only 14.25% (722) had a shared component. This is consistent with complex cell type heterogeneity in bulk tissues [27] since there is more power to discover eQTLs with pleiotropy across the underlying cell types.

### CONTENT more accurately distinguishes disease-relevant genes than traditional TWAS approaches in simulated data

We performed a simulation study in which we evaluated the sensitivity, specificity, and power of CONTENT, UTMOST, and context-by-context to implicate the correct gene in TWAS. In our experiments, we simulated a phenotype in which 20% of the variability was composed of the genetically regulated expression of 300 randomly selected gene-context pairs (100 genes and 3 contexts each). We simulated gene expression for 1,000 genes across 20 contexts as before, however, to capture a range of genetic architectures in the simulation, for each gene, we sampled from a standard uniform distribution to determine the proportion of shared variability. We varied the heritability of gene expression and considered power as a method’s ability to discover the correct genes associated with a phenotype. To compare power, we calculated the area under receiver-operating curve (AUC) using the maximum association statistic for a given gene across contexts.

Across simulations, CONTENT(Full) was the highest powered in terms of gene discovery (Figure 5). CONTENT(Shared) performed very similarly to CONTENT(Full) in the setting with the lowest heritability, however, our simulations show the necessity for CONTENT(Full) as it substantially outperforms both CONTENT(Specific) and CONTENT(Shared) across a range of heritabilities. Moreover, CONTENT(Full) significantly outperformed both the context-by-context approach and UTMOST. Specifically, the range of percent change in AUC of CONTENT(Full) over previous methods was as follows: CONTENT(Shared) 1.9%-9.9%; CONTENT(Specific) 13.6%-22.4%; UTMOST 2.2%-8.6%; context-by-context 1.2%-10.6%. Generally, we observed that CONTENT(Full) was its most powerful for genes in which there was both shared and specific effects, UTMOST was its most powerful in settings with high sharing, and the context-by-context approach was its most powerful in settings with low sharing and high specificity of genetic effects within contexts.

**Figure 5.**
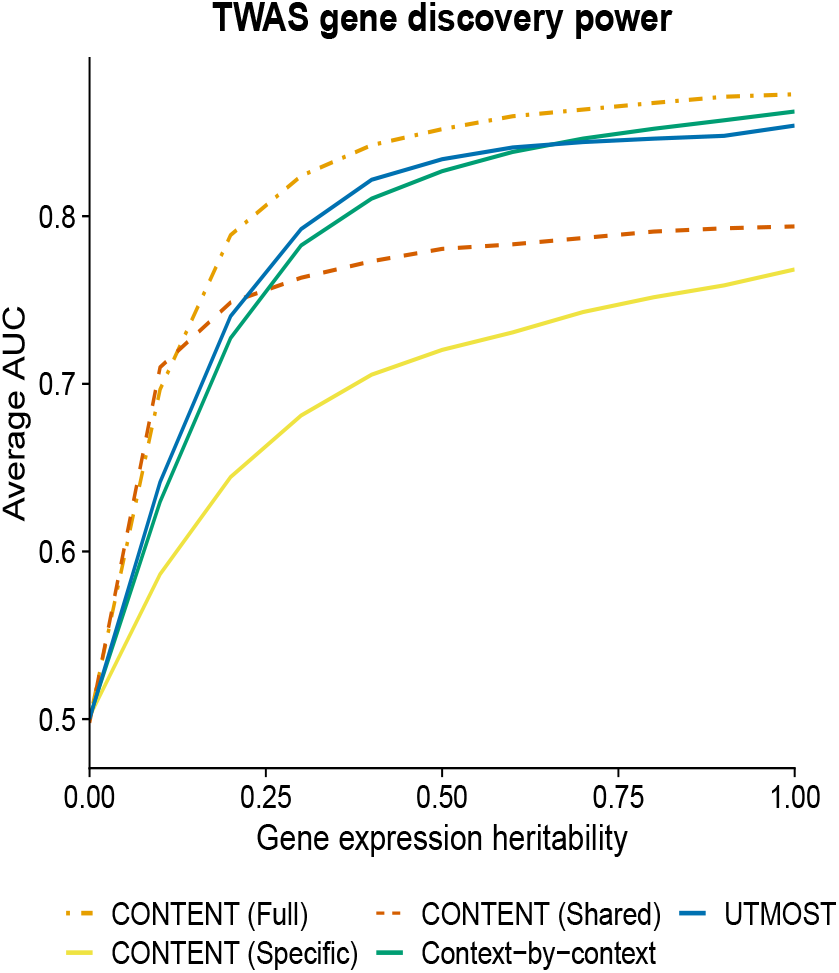
CONTENT(Full) is powerful, sensitive, and specific in simulated TWAS data. Average AUC from 1,000 TWAS simulations while varying the overall heritability of gene expression. Each phenotype (1,000 per proportion of heritability) was generated from 300 (100 genes and 3 contexts each) randomly selected gene-context pairs’ genetically regulated gene expression, and the 300 gene-context pairs’ genetically regulated expression accounted for 20% of the variability in the phenotype. In genes with low heritability, CONTENT(Shared) performed similarly to CONTENT (Full), however CONTENT(Full) was the most powerful method in discovering the correct genes for TWAS across the range of heritability. CONTENT(Full) was significantly more powerful than UTMOST and the context-by-context approach at all levels of heritability.

As with previous methods [9], we performed simulations in which the causal context(s) has been observed. In real data applications, this may not occur, and in such cases, further complexities may arise due to genetic correlation. We report a brief set of experiments evaluating fine-mapping gene-context pairs in TWAS when all contexts are observed (see Supplementary; Figures S12, S13), however, the complexities posed by missing tissues and cell types are beyond the scope of this manuscript, and we therefore leave the development of relevant methodology as future work.

### Application of CONTENT to TWAS yields novel discoveries over previous methods

We performed TWAS across 22 complex traits and diseases collected from a variety of GWAS [28–41] using weights trained by CONTENT, UTMOST and the context-by-context approach on GTEx and CLUES. We passed forward weights to FUSION-TWAS[3]—a software that performs TWAS using GWAS summary statistics and user-specified gene expression weights—for a gene-context pair if the pair’s expression was predicted at a nominal p-value less than .1 (See Methods; Figure S14).

Across all traits at an hFDR of 5%, CONTENT discovered 47% and 234% more associations (unique TWAS genes) than the context-by-context approach and UTMOST respectively with GTEx weights, and 160% and 459% more associations than the context-by-context approach and UTMOST respectively with weights built from the CLUES dataset (Table 1). We find that, with GTEx weights, the associations implicated by the context-by-context approach had more overlap with the associations implicated by CONTENT(Specific) (median Jaccard similarity (JS) across traits =.406) than CONTENT(Shared) (JS=.177). This is consistent with our simulation results in which the context-by-context approach was most powerful in cases of high context-specificity and low context-sharing (Figures S12, S13). The associations discovered by UTMOST, which leverages pleiotropy, had similar overlap with CONTENT(Shared) (JS=.175) as well as CONTENT(Specific) (JS=.182). With CLUES weights, the context-by-context approach again had greater similarity with CONTENT(Specific) (JS=.242) than CONTENT(Shared) (JS=.078), whereas UTMOST discovered TWAS genes that overlapped more with CONTENT(Shared) (JS=.100) than CONTENT(Specific) (JS=.085). As UTMOST, CONTENT, and the context-by-context approach discovered both overlapping and unique associations, we suggest that the approaches complement—rather than replace—one another.

**Table 1.** CONTENT outperforms existing methods in TWAS across 22 complex traits and diseases. TWAS results (unique loci, merging genes within 1MB) across 22 complex traits and diseases using weights output by CONTENT, UTMOST, and the context-by-context method. CONTENT(All) refers to the collection of all loci output by at least one CONTENT model. CONTENT(Full) added an average of 15% and 19% of gene-trait discoveries over the CONTENT(Shared) and CONTENT(Specific) approaches together at an hFDR of 5% in GTEx and CLUES respectively. See Supplementary Table S1 for GWAS trait information.

| Trait | GTEx |  |  |  |  |  | CLUES |  |  |  |  |  |
| --- | --- | --- | --- | --- | --- | --- | --- | --- | --- | --- | --- | --- |
|  | Tissue-by-tissue | UTMOST | CONTENT (All) | CONTENT (Full) | CONTENT (Specific) | CONTENT (Shared) | Tissue-by-tissue | UTMOST | CONTENT (All) | CONTENT (Full) | CONTENT (Specific) | CONTENT (Shared) |
| AD | 18 | 14 | 25 | 20 | 14 | 10 | 8 | 4 | 10 | 6 | 9 | 2 |
| Asthma | 145 | 135 | 229 | 181 | 185 | 81 | 57 | 52 | 101 | 75 | 88 | 19 |
| Bipolar | 34 | 59 | 83 | 68 | 59 | 30 | 3 | 12 | 31 | 21 | 24 | 8 |
| CAD | 8 | 12 | 16 | 13 | 13 | 4 | 6 | 2 | 8 | 6 | 5 | 2 |
| CKD | 24 | 24 | 39 | 28 | 28 | 15 | 5 | 6 | 14 | 8 | 12 | 1 |
| Crohn's | 80 | 66 | 104 | 74 | 92 | 42 | 30 | 28 | 54 | 45 | 39 | 12 |
| Eczema | 21 | 19 | 42 | 35 | 28 | 10 | 9 | 2 | 18 | 14 | 13 | 1 |
| FastGlu | 19 | 15 | 16 | 13 | 14 | 7 | 7 | 6 | 10 | 7 | 10 | 3 |
| HDL | 58 | 51 | 83 | 70 | 72 | 38 | 24 | 14 | 28 | 23 | 24 | 8 |
| IBS | 2 | 7 | 8 | 8 | 3 | 1 | 1 | 1 | 1 | 1 | 1 | 1 |
| LDL | 94 | 72 | 125 | 111 | 104 | 62 | 28 | 21 | 62 | 49 | 50 | 13 |
| Lupus | 100 | 66 | 148 | 107 | 118 | 46 | 32 | 19 | 59 | 43 | 44 | 12 |
| MDD | 92 | 93 | 150 | 111 | 121 | 55 | 28 | 20 | 46 | 33 | 35 | 11 |
| MS | 17 | 10 | 31 | 28 | 18 | 5 | 6 | 4 | 12 | 9 | 9 | 4 |
| PBC | 52 | 39 | 66 | 57 | 56 | 29 | 18 | 20 | 21 | 16 | 18 | 3 |
| Psoriasis | 35 | 25 | 54 | 41 | 38 | 15 | 16 | 9 | 23 | 18 | 20 | 6 |
| RA | 85 | 61 | 102 | 81 | 86 | 40 | 41 | 24 | 47 | 34 | 39 | 11 |
| Sarcoidosis | 8 | 12 | 24 | 17 | 16 | 7 | 4 | 3 | 7 | 6 | 5 | 1 |
| Sjogren | 9 | 8 | 9 | 8 | 6 | 2 | 2 | 1 | 5 | 3 | 5 | 1 |
| T1D | 78 | 57 | 109 | 82 | 97 | 40 | 33 | 25 | 54 | 41 | 44 | 14 |
| T2D | 170 | 144 | 231 | 203 | 191 | 108 | 59 | 43 | 82 | 59 | 71 | 15 |
| Ulc colitis | 9 | 8 | 16 | 13 | 10 | 3 | 1 | 2 | 9 | 8 | 8 | 1 |

We next compared the different CONTENT models to understand their properties in real data. With GTEx weights, CONTENT(Full) replicated an average of 99.0% and 66.8% of the associations discovered by CONTENT(Shared) and CONTENT(Specific) respectively (hFDR ≤ 5%). CONTENT(Full) replicated an average of 78.4% and 63.9% of the associations discovered by CONTENT(Shared) and CONTENT(Specific) respectively with the CLUES weights. Notably, CONTENT(Full) is the best predictor out of all the CONTENT models on average, and particularly when there exist both shared and specific effects. Consequently, across all traits, the inclusion of CONTENT(Full) with CONTENT(Shared) and CONTENT(Specific) led to an average increase of 15% and 19% in the number of genes with significant TWAS associations with GTEx weights and CLUES weights respectively.

We investigated the genes implicated by CONTENT(Full) that were not significant in CONTENT(Shared) or CONTENT(Specific) and found that many of the discoveries replicated known gene-trait associations. Using weights built from GTEx for example, CONTENT(Full) discovered a significant association of coronary artery disease (CAD) and VEGFC (p=8.67e-07, artery aorta), a gene whose serum levels have been significantly associated with cardiovascular outcomes [42]. Furthermore, CETP, which is thought to be involved in atherogenesis and HDL levels [43, 44], was not implicated by either CONTENT(Shared) or CONTENT(Specific), but was implicated in the TWAS of HDL with CONTENT(Full) (p=1.26e-170, whole blood). CONTENT(Full) also discovered a significant association of myelin oligodendrocyte glycoprotein (MOG) and rheumatoid arthritis (RA) (minimum p value across tissues; p=2.55e-11, brain amygdala), whereas CONTENT(Shared) and CONTENT(Specific) did not. RA patients have been shown to have significantly higher levels of anti-MOG IgG than controls [45].

Using weights built from CLUES, CONTENT(Full) also led to increased power over CONTENT(Shared) and CONTENT(Specific). Namely, CONTENT(Full) replicated an association between asthma and RHOA (B cell, p=1.22e-04), which is involved in smooth airway contraction possibly through inflammation [46, 47]. Moreover, we also discovered a significant association of Type 2 Diabetes (T2D) and SIRT5 (CD4 T cell, p=2.23e-04) using CONTENT(Full), and note that SIRT5 has previously been indicated to play roles in metabolism and beta-cell functionality [48, 49]. Finally, CONTENT(Full) indicated an association of APH1A with eczema (conventional dendritic cell, p=3.49e-07). APH1A is involved with Notch signaling, which when disrupted in the skin, contributes to abnormalities such as eczema [50].

Moreover, the genes implicated by CONTENT but neither UTMOST nor the context-by-context approach replicated previously associated genes-trait pairs, several of which with known biological relationships to the trait of interest. Within Alzheimer’s disease, these genes included CBLC[51, 52], MS4A4A [53], and MADD [54] with the GTEx weights, as well as VASP[55, 56], RELB[51], TRAPPC6A[57, 58] with CLUES weights. Additionally, in Crohn’s disease, CONTENT implicated the following genes, whereas previous methods did not: LRRC26[59] and RASSF1A[60] with GTEx weights, as well as CARD6 (an inhibitor of NOD)[61, 62] using CLUES weights. For major depression disorder (MDD), CONTENT implicated CAMP[63] using GTEx weights, and FLOT1 [64, 65] using CLUES weights.

## 3 Discussion

In this work, we introduce CONTENT, a computationally efficient and powerful method to estimate the genetic contribution to expression in multi-context studies. CONTENT can distinguish the context-shared and context-specific components of genetic variability and can account for correlated intra-individual noise across contexts. Using a range of simulation and real studies, we showed that CONTENT outperforms previous methods in terms of prediction accuracy of the total genetic contribution to expression variability in each context. Interestingly, we also found that when there exists a gene with a genetic component of expression, the heritability is often dominated by the context-specific effects at the single-cell level, but at the tissue level, the heritability is dominated by the context-shared effects. Finally, CONTENT was more powerful, specific, and sensitive than previous approaches in applications to TWAS.

Using weights trained by CONTENT, UTMOST and the context-by-context approach, we discovered 12,150 unique gene-trait associations through TWAS. To our knowledge, we present the first application of TWAS trained on a single-cell RNAseq dataset for a collection of 90 individuals’ PBMCs. For both the weights generated by GTEx and CLUES, CONTENT was largely more powerful than UTMOST and the context-by-context approach in TWAS. However, we emphasize that the approaches often capture genes unique to each approach. Each method may therefore complement each other and may be combined in TWAS to maximize the number of discoveries made as different methods are likely favorable under different genetic architectures. Though we show that CONTENT may be useful in fine-mapping the specific tissue relevant for a TWAS association in simulations, we note that fine-mapping to the correct tissue in real data is a particularly difficult task. For example, throughout this manuscript, we assume that the causal tissue is included in the measured tissues, however, when this is not the case, CONTENT and all TWAS approaches may associate an incorrect, correlated tissue. For example, in the case of chronic kidney disease, CONTENT implicated GATM–a gene thought to be involved with kidney disease and GFR levels [66–68]–however, there were significant associations with many tissues including the tissue-shared component. This may be due to the fact that kidney expression is not measured in this version of the GTEx dataset. Future work may explore using the CONTENT-trained weights and jointly fitting all TWAS Z scores, or otherwise accounting for missingness.

We also leveraged recently developed methodology for controlling the false discovery rate when summarizing significantly predicted genes, gene-contexts, and TWAS associations [17, 18]. This approach has been shown to effectively control the FDR across contexts in eQTL studies, and to our knowledge, it is the first time such an approach has been used to effectively control the FDR when predicting expression values and when making discoveries using TWAS. While our analyses focused on the comparison of CONTENT, UTMOST, and the context-by-context approach, we emphasize that by using this type of false discovery correction, all methods can be used in combination with one another, rather than in replacement of one another. For downstream analysis, combining all prediction methods is crucial, as certain genes or gene-context pairs may be (better) predicted by one method and not others. In the GTEx data for example, when we included models built by UTMOST and the context-by-context approach to the correction scheme for CONTENT, the number of genes for which there was a significant model for a given tissue increased on average by 7.56%.

Importantly, neither UTMOST nor the context-by-context method distinguishes the context-specific and context-shared components of genetic effects on expression. Implicitly, by modeling all contexts independently, the context-by-context fit is best-suited for cases in which there is no effect-sharing across contexts. As UTMOST considers all contexts simultaneously, its power is maximized in cases where the genetic effects are mostly shared. Additionally, these methods do not account for the shared correlated residuals between samples, thus they do not maximize their predictive power.

While a previous approach proposed by Wheeler et al. [14] does model the correlated intra-individual noise, CONTENT offers several advantages. The previous decomposition does not include an option to leverage both the context-shared and context-specific components of expression to form a final predictor of the observed expression for a given context. We show that this is especially crucial in the context of single-cell data wherein the prediction accuracy for a given gene-context increases drastically when using both components (Figure 3). Further, without properly combining both components (e.g. via regression), the context-specific genotype-expression weights produced by the previous decomposition may have the incorrect sign, as they are considered residuals of the context-shared component and are not properly re-calibrated to the observed expression. Unlike the novel decomposition proposed by CONTENT, this previous approach also does not intuitively allow for additional sources of pleiotropy or effects-sharing (see Supplementary Text for discussion of brain level sharing in GTEx). Finally, the decomposition used in the previous method is based on a linear mixed model fit on a per-gene basis, and is therefore much less computationally efficient.

In this manuscript we focused on prediction of the total genetic contribution to expression as well as the context-shared and context-specific components of expression. Nonetheless, future work using the methodology presented here can be extended to a wide variety of problems. Primarily, the decomposition can be used to efficiently estimate Gene×Context heritability using existing software for heritability estimation, e.g. *GCTA* [69], on the decomposed components offering computational speed up over existing methods for cross-context heritability estimation [26]. Additionally, the decomposed components from CONTENT may also be included in previous approaches, e.g. UTMOST, to gain further power. Further, by training each method on the single-cell level data, we offer researchers the means to pursue their own association analyses at a lower level of granularity than was previously available. The finding that single-cell data may have lower levels of effects-sharing than tissue-level data may also spark investigations into the biological mechanisms (e.g. more specific regulation) and statistical mechanisms (e.g. sample heterogeneity confounding) by which this can occur.

In conclusion, the increased prediction accuracy, specificity, computational speed, and hierarchical testing framework of CONTENT will be paramount to unveiling context-specific effects on disease as well as uncovering the mechanisms of context-specific genetic regulation.

## Code and data availability

Trained weights for the GTEx V7 dataset and our in-house single-cell RNAseq are available at TWAShub (http://twas-hub.org/). The CONTENT software is freely available at https://github.com/cozygene/CONTENT. We provide TWAS summary statistics for all three methods on both datasets (as well as an indicator of whether the association was hierarchical FDR-adjusted significant) at doi.org/10.5281/zenodo.5209239.

## Author contributions

NZ and BB conceived of the project and developed the statistical methods with MT. MT implemented the comparisons with simulated data with contributions from AT. MT, AL, and MGG, performed the analyses of the GTEx and CLUES data and additional analyses. MT implemented the software. MT, NZ, and BB wrote the manuscript, with significant input from EH, CJY, AG, MGG. AG prepared the online data resources.

## Conflicts of interest

CJY is a Scientific Advisory Board member for and holds equity in Related Sciences and ImmunAI. CJY is a consultant for and holds equity in Maze Therapeutics. CJY is a consultant for TReX Bio. CJY has received research support from Chan Zuckerberg Initiative, Chan Zuckerberg Biohub, and Genentech.

## 4 Methods

### An overview of the CONTENT model

In this section, we detail the assumed generative model and objectives of CONTENT. CONTENT is based on the methodology and decomposition of a previous work by Lu et al., FastGxC [16]. In brief, like FastGxC, we assume that the expression of an individual in a given gene and context is a combination of a context-shared genetic component that is shared across different contexts and a context-specific genetic component that is specific to a context, that is

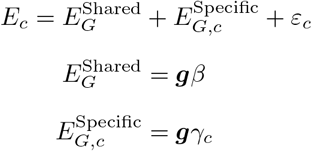

where *E_c_* denotes the expression of the individual at the gene in context *c*, 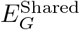 and 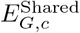 denote the components of the expression due to context-shared and context-specific genetic effects respectively, *β* and *γ_c_* represent the context-shared and context-specific cis-genetic effects respectively, ***g*** the individual’s cis-genotypes and 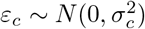 represents the environmental effects (and non-cis-genetic effects) on the individual’s gene expression.

The objective of CONTENT is to build a genetic predictor of context-specific phenotypes. While previous work has focused on building powerful genetic models for *E_c_*, we aim to build unbiased models that partition and estimate the context-shared ***g****β* and context-specific terms ***g****γ_t_*. Specifically, we aim to maximize the power to detect the context-specific terms, allowing some leniency in the accuracy of context-shared terms, as we are interested in context-specific effects. Moreover, as a context-specific predictor can be used in downstream analyses to identify the specific context(s) through which genetic variation manifests its effect on the phenotype and disease risk, we also aim to minimize the correlation between the predicted context-specific component and the true context-shared component. Finally, our method must account for the correlated intra-individual noise across contexts, and do so in a computationally efficient manner.

### Decomposing multilevel data

Many genomic datasets, such as those of GTEx, have a multilevel nature; first the individuals are sampled, and second an individual is measured in each context. To take the multilevel structure of the data into account, the observed expression on gene *j* can be decomposed into an offset term, a between-individual component and a within-individual component [70]. That is, if *E_ijc_* denotes the observed expression level for individual *i* (*i* = 1, …, *I*) on gene *j* (*j* = 1, …, *J*) and context *c* (*c* = 1, …, *C*), *E_ijc_* can be decomposed as

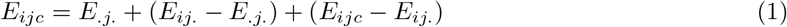

where 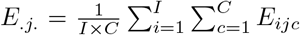 the mean expression of gene *j* computed over all (*I*) individuals and all (*C*) contexts, and 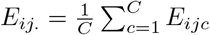 the mean expression of individual *i* on gene *j*, computed over all contexts. In (1), *E_.j._* is a term that is constant across individuals and contexts for each gene, (*E_ij._* − *E_.j._*) is the between-individuals deviation, and (*E*_*ijc*_ − *E*_*ij*._) is the within-individual deviation of the expression on gene *j* in context *c*.

Variables that differ between but not within individuals, e.g. sex and genotype, will have an effect on (*E*_*ij*._ − *E*_.*j*._) but not on (*E*_*ijc*_ − *E*_*ij*._). On the other hand, variables that change within individuals but are the same between individuals, e.g. the genetic effect on a specific context, will have an effect on (*E*_*ijc*_ − *E*_*ij*._) but not on (*E*_*ij*._ − *E*_.*j*._).

In the context of estimation, we first center and scale the expression of gene *j* in each context *c*, i.e. 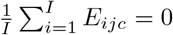 and 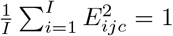. Therefore, 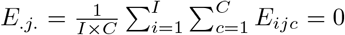, and equation (1) simplifies to:

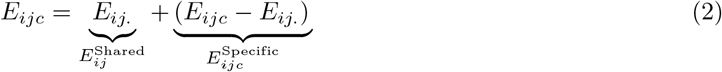

### A formal description of CONTENT

We use the simplified decomposition in equation (5) to build genetic predictors of context-specific effects while accounting for the correlated intra-individual noise across contexts. Intuitively, the between-individuals variability serves as the component of expression that is shared across contexts, *E*^Shared^, and the deviance from this shared component (i.e. the within-individual variability) serves as the context-specific component of expression, *E*^Specific^. Moreover, treating the context-specific component as a deviance from the context-shared component leads the decomposition to have the property that as the correlation of intra-individual noise across contexts increases, the power to detect context-specificity also increases. In addition, the decomposition generates context-shared and context-specific components of expression that are orthogonal to each other. Further rationale for using the decomposed expression is included Supplementary Section 1 and the text by Lu et al. [16]. Lu et al. also include a description of the decomposition’s equivalence to a linear mixed model.

For a single gene *j*, CONTENT takes as input centered, scaled, and residualized (over a set of covariates) expression measured across *I* individuals in *C* contexts and an *I* × *m* genotype matrix *G_j_* with *m* measured cis-SNPs for gene *j*. CONTENT then decomposes the expression vectors into *C* context-specific components and a single context-shared component by simply calculating the mean of expression for each individual across contexts, and setting the context-specific expression for context *c* as the difference between the observed expression of context *c* and the calculated context-shared expression. As it has been observed that cis-genetic effects may be sparse and that the elastic net may perform best relative to other penalized linear models in the context of genetically regulated gene-expression [4, 14], CONTENT fits *C* + 1 penalized linear models for the *C* + 1 expression components using an elastic net. Lastly, CONTENT generates a final genetic predictor of expression by combining the context-shared and context-specific components. Importantly, as the context-specific component is a deviance from the context-shared component, the sign of the context-specific component must be properly realigned when combining both components of expression to make a final predictor. We refer to this linear combination of expression components as the “full” model of CONTENT and fit it using a simple linear regression:

1. Obtain 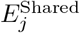 and 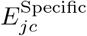 from the decomposition.
2. Generate cis-genetic predictors of each component using cross-validated elastic net:

a. Fit cross-validated elastic net regressions for the shared and specific components:

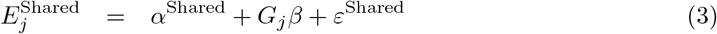

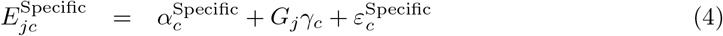
b. Use the estimates to generate genetic predictors of each component:

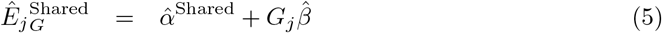

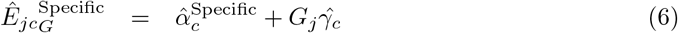
3. Regress the expression of context *c* onto the context-shared and context-specific components:

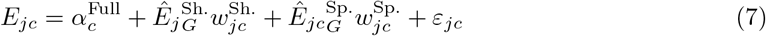

Within each regression, *α* represents the offset and we assume that all *ɛ* are from a normal distribution with mean 0 and standard deviation that is a function of the given outcome.

We save for each gene the set of estimated regression weights 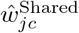 and 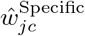 from equation (4) for use in downstream analyses. Namely, in TWAS, each context receives a single vector of weights, and to test the association of a gene-context’s full model to a trait, we simply use a weighted sum of the predictors learned from equation (3), 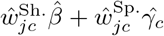. We also use the same procedure for the context-specific weight to ensure the correct directionality. To test for significance of genetic effects (i.e. to call an eGene or eAssociation), we correlate each component of expression—the context-shared, context-specific, and full—to its corresponding genetically predicted value.

### Controlling the false discovery rate across contexts

Generally, methods for building genetic predictors of expression or TWAS predictors leverage either Bonferroni correction or false discovery rate (FDR). Nonetheless, using a Bonferroni correction may be too stringent (for example, as tests across contexts may be correlated), and using FDR within each context or across all contexts simultaneously may lead to an inflation or deflation to the false disovery proportion within certain contexts [17]. To simultaneously control the FDR across all contexts at once, a hierarchical false discovery correction— treeQTL—was developed [17]. The treeQTL procedure leverages the hierarchical structure of a collection of tests (e.g. gene level and gene-context level) to properly control the FDR across an arbitrary number of contexts and levels in the hierarchy as well as boost power in cases where a gene has a significant association in multiple contexts [6, 17, 18]. (See Supplementary Methods for further intuition.)

Notably, using CONTENT, our testing hierarchy contains 3 levels; (1) at the level of the gene, (2) at the level of the context, and (3) at the level of the method or model (Figure S3). Intuitively, a gene may contain a genetic component that is shared across all contexts, or a given context may have its own genetic architecture. In CONTENT, a given context may have its own genetic predictor from either the context-specific component or the full model. Using treeQTL with this structure is robust across multiple contexts, and since the tree is structured such that a specific method/model is at the final level of testing for a context, it enables incorporation of additional models trained from other approaches (such as those fit on a context-by-context basis or by UTMOST). Moreover, we can add to the shared leaf an additional level of tests to account for additional components of effects-sharing, such as a brain tissue-shared component.

### Comparison to other methods

We compared the prediction accuracy of CONTENT to a context-by-context TWAS model [3, 4] in which the expression of each context is modeled separately, and to UTMOST [9], a method that jointly learns the genetic effects on all contexts simultaneously. Specifically the model based on TWAS fits a penalized linear model for each context. UTMOST, on the other hand, employs a group LASSO penalty across all contexts simultaneously, allowing it to gain power over the context-by-context approach by considering all individuals and contexts in a study at once. As we were we able to use a fast R package for penalized regression[71], we used 10-fold cross-validation to fit the context-by-context model. Owing to UTMOST’s computational intensity, we used its default value of 5 folds for cross-validation.

We also compared CONTENT to a previous approach by Wheeler et al., orthogonal tissue decomposition (OTD)[14]. OTD is a direct correlate of CONTENT(Shared) and CONTENT(Specific), and is generated by fitting a mixed effects model across all contexts for a given individual. Namely, a mixed effects model is fit as follows: an individual’s expression across all tissues is set as the outcome, the shared expression is modeled as a random individual-level intercept and is estimated using the posterior mean, and the specific expression is treated as the residuals from the fit model (after adjusting for covariates). Under infinite sample sizes, the components of OTD are equivalent to CONTENT(Shared) and CONTENT(Specific).

### Evaluations on GTEx and CLUES

We residualized the expression of each gene in each context over their corresponding covariates (e.g. PEER factors, age, sex, batch information) prior to fitting UTMOST and an elastic-net model for each context in the context-by-context approach. We did the same residualization before decomposing and then fitting the context-shared and context-specific components with an elastic net for CONTENT. After generating cross-validated predictors for each method, we examined the number of significantly predicted genes as well as the prediction accuracy (in terms of adjusted *R*^2^) between the cross-validation-predicted and true gene expression per gene-context pair.

To properly control the false discovery proportion at .05 across-contexts and within-methods, we employed a hierarchical FDR correction [17, 18] separately for CONTENT, UTMOST, and the context-by-context approaches. Notably, using this correction for all methods provides a generous comparison to previous methods, as when there exists at least one significantly heritable gene-context association for a given gene, there is a relative gain in power over the context-by-context FDR for other contexts tested within this gene [17, 18].

### Application to TWAS

We performed transcription-wide association studies across 24 phenotypes using FUSION-TWAS[3]. FUSION-TWAS uses GWAS summary statistics and user-specified gene expression weights with an LD reference panel to perform the test of association between genetically predicted gene expression and a phenotype of interest. We tested a gene-context pair for association if the pair’s expression was predicted at a nominal p-value of .1, and note that this threshold does not substantially alter the number of TWAS discoveries (Figure S14). Notably, previous methods may use their own test of gene-context-trait association or leverage set tests (e.g. Berk Jones[9]) to combine their associations across all contexts for a given gene and therefore increase power. In this comparison, we report the association as output by FUSION (a single gene-context-trait association) and corrected by hierarchical false discovery without any sort of set test for the sake of equality in the comparison. We ran FUSION-TWAS using the default recommended settings, with reference data from the 1000 genomes project [72]. TWAS weights were trained on the GTEx v7 dataset[2] as well as the CLUES[20] single-cell RNAseq dataset of PBMCs.

### Simulations to evaluate prediction accuracy

To evaluate the properties of our method relative to other methods we perform a series of simulation experiments. We first simulate genotypes for each individual, where each individual *i* and each locus *m* (*m* = 1 : *M*) is independent, and there are no rare SNPs:

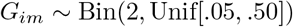

We then draw both context-shared (*β*_*j*._) and context-specific (*β*_*jc*_) effect sizes for each SNP from a normal distribution with a Bernoulli random variable *I*_*m*_ controlling the probability that the *m*^*th*^ SNP is causal (i.e. induce sparsity of genetic effects).:

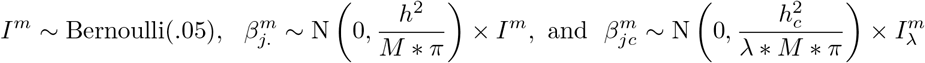

Here, *h*^2^ and 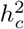 are the context-shared and context-specific heritabilities of expression on gene *j*. In general, the SNPs with nonzero context-specific effect sizes were subsampled from SNPs with nonzero context-shared effect sizes. We additionally simulate for a subset of contexts some number of truly context-specific eQTLs drawn from Poisson(*λ* = 1) for randomly selected SNPs that were not eQTLs for the context-shared effects. Finally, we simulate the expression of gene *j* as follows:

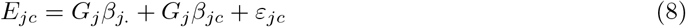

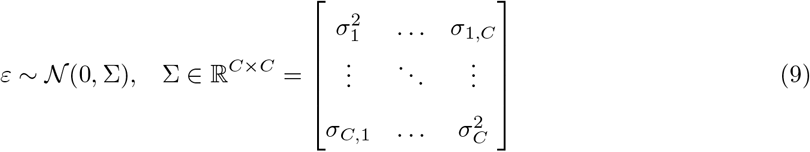

where *ɛ* ∈ ℝ^*I*^, represents the correlation of environment or intra-individual noise across contexts, 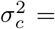 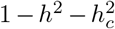 is the variances of each context *c*, and 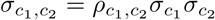 is the covariance of context *c*_1_ and *c*_2_. We generated data under varying levels of context-specific heritability, truly context-specific eQTLs, causal SNPs, and correlation of intra-individual noise across contexts. The number of contexts was set to 20, and to replicate a setting similar to GTEx, the corresponding sample sizes of each ranged from 75 to 410 where individuals were not necessarily measured in every context. In our simulations, we generated one train and one test data set using the above framework. We evaluated the performance of each method by comparing the true and predicted expression in the test data set, using the predictor learned from the training data set.

To assess the effect of additional sharing on a subset of contexts, we also set up a simulation framework using the same generative process as above, only that a subset of contexts also received additional genetic effects. More rigorously, for this subset of contexts (acting as brain contexts in GTEx, for example), expression was generated as in equation (6) with an additional term:

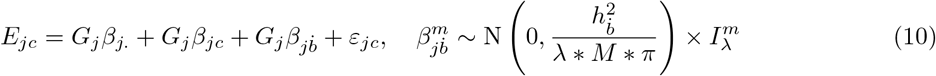

where each variable is simulated as before, 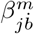 corresponds to additional genetic effects that are subsampled from SNPs that have a context-shared effect, and 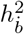 is the brain-shared heritability.

### Simulations of TWAS performance

Using the above generated genotypes and gene expression, we simulated phenotypes to evaluate the performance of each method under the assumed model in TWAS. For a given phenotype, we randomly selected 300 gene-context pairs (100 genes, 3 contexts each) whose expression would comprise a portion of a phenotype. Explicitly, we generated a phenotype as follows:

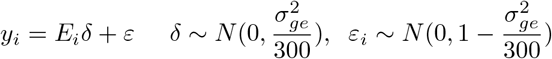

Where *E_i_* is the standardized genetic expression of the 300 gene-context pairs for individual *i*, *δ* is the length-300 vector of effect sizes for each gene-contexts’ expression, 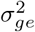 is the variance in the phenotype *y_i_* due to cis-genetic gene expression, and *ɛ*_*i*_ corresponds to environmental effects (or noise) as well as trans-genetic effects for individual *i*. In our simulations, we varied the heritability of gene expression and fixed variability in the phenotype due to genetic gene expression to .2. To simulate a wide range of genetic architectures, the proportion of heritability of gene expression due to the context-shared effects was sampled from a standard uniform distribution, and the proportion of heritability due to context-specific effects was (1-the context-shared proportion). Once we generated a phenotype, we performed a TWAS using weights output from each method by imputing expression into a simulated external, independent set of 10000 genotypes that followed the same generation process as in the previous subsection.

## Supplementary Methods and Information

### Intuition for using the decomposition to model genomic features

The decomposition described in the methods section lays a framework for CONTENT as it directly accounts for the shared noise and generates orthogonal context-shared and context-specific components of genomic features. First, we note that in multi-context data, repeated measurements of one individual will likely have correlated errors; in the context of GTEx data, an individual’s environment as well as technical noise is likely to affect their expression in all contexts. The above decomposition exploits this structure, which improves the power to learn the context-specific variability of expression. Put more rigorously, consider the expression of gene *j* in an individual measured in a baseline context and then again after a stimulation:

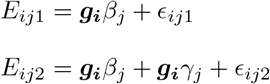

Where *E*_*ij*1_ and *E*_*ij*2_ denote the observed expression level of individual *i* at gene *j* at baseline and stimulation respectively, ***g_i_*** represents a vector of the individuals’ genotype at some nearby *cis*-SNPs, *β*_*j*_ denotes the baseline genetic effects on expression, *γ*_*j*_ denotes the stimulation-related genetic effects on expression, and *ϵ*_*ij*1_ and *ϵ*_*ij*2_ represent the environmental effects (or noise) on the individual’s expression of gene *j* in baseline and stimulation respectively. In teasing apart the genetic effects that are different after stimulation, one might examine the difference in the expression between contexts:

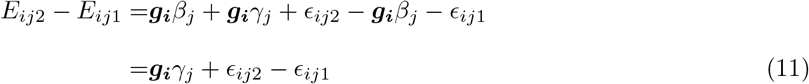

which leaves only the difference in expression due to the stimulation-specific, or in other words, context-specific component, and noise. Under the scenario in which the errors are perfectly correlated, (11) simplifies to:

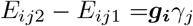

Clearly, this will greatly increase our ability to build a genetic model of the stimulation-specific component. In terms of CONTENT, the baseline genetic effects correspond to the context-shared genetic effects, and the stimulation-specific effects correspond to the context-specific effects. Put simply, we propose the context-shared genetic effects be considered a “baseline” effect, and that the context-shared genetic effects are simply offsets to the context-shared effect. This model is directly related to equation (3):

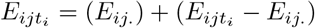

where *E*_*ij*._ and 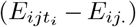 correspond to the context-shared and context-specific genetic effects respec-tively. By construction, *E*_*ij*._ and 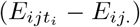 are orthogonal, and thus we have generated orthogonal components for the context-shared and context-specific components of expression.

**Figure S1.**
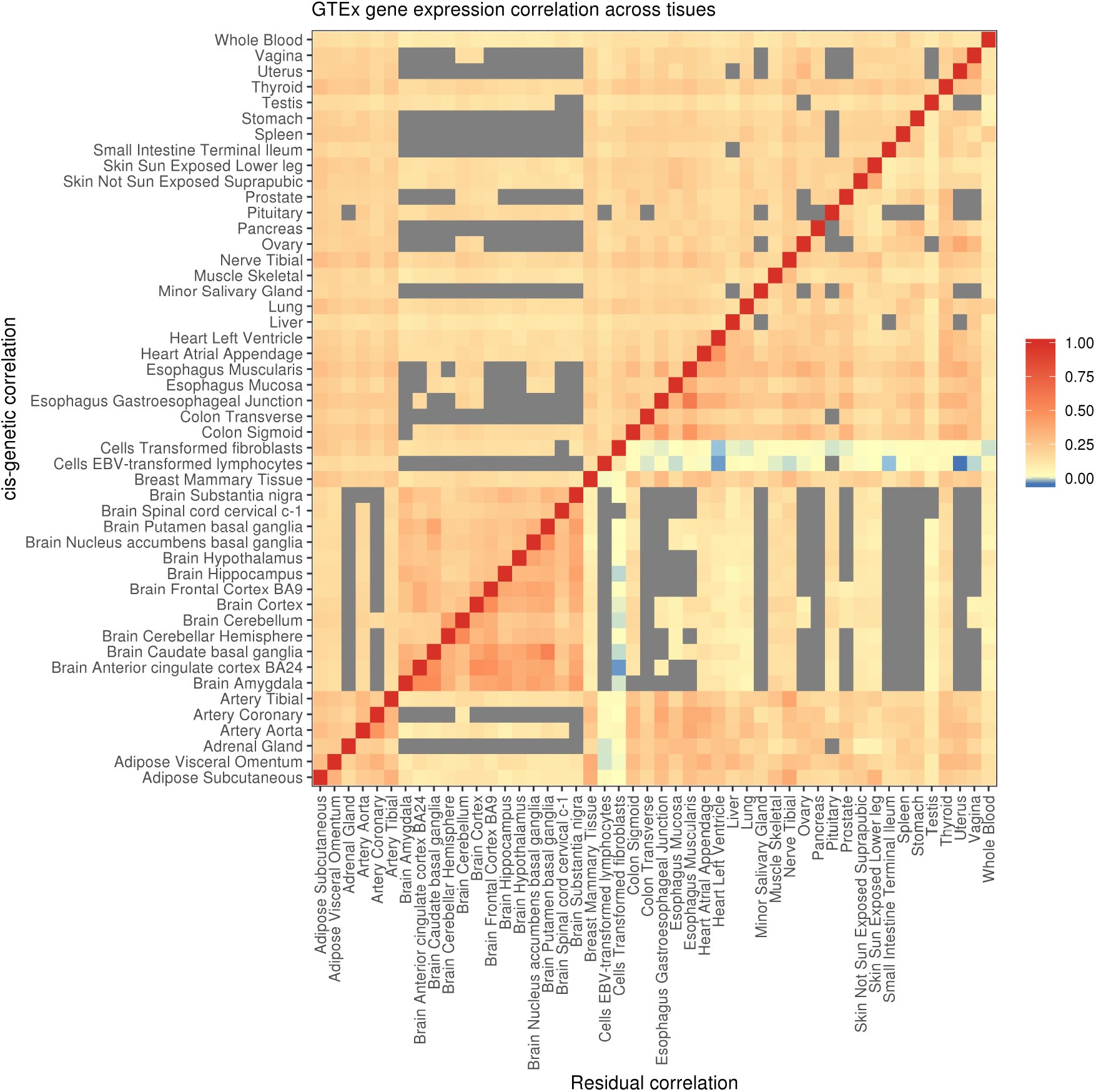
Gene expression correlation across tissues in the GTEx study. Using a linear mixed model with bivariate REML [69, 73], we calculated cis-genetic and residual (which captures variance due to both trans-genetic effects as well as residual effects) variance and covariance components for each gene-tissue pair across GTEx. The gray units indicate tissue pairs with less than 10% sample overlap. In both the genetic (upper) and residual (lower) components, there was widespread cis-genetic and residual correlation, with the brain tissues showing higher correlations compared to other tissues.

**Figure S2.**
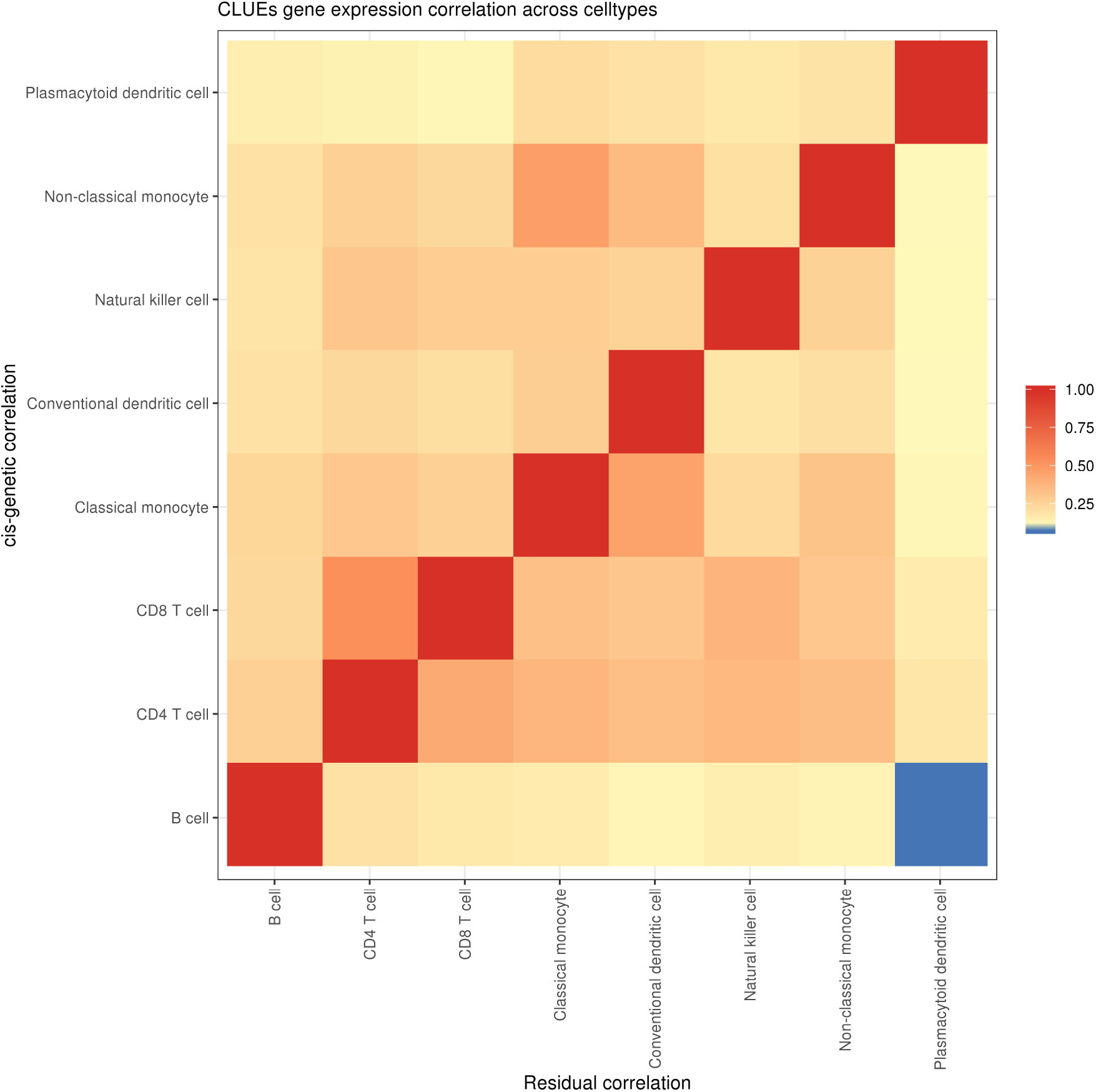
Gene expression correlation across cell types in the CLUEs study. Using a linear mixed model with bivariate REML[69, 73], we calculated cis-genetic and residual (which includes trans-genetic effects) variance and covariance components for each gene-cell type pair across CLUEs.

### Hierarchical false discover correction

Multiple hypotheses correction in the context of discovering genes, gene-context pairs, and downstream associations of genetically-regulated gene expression with phenotypes varies across approaches [3, 4, 15]. For discovering gene and gene-context associations, previous approaches often leverage a Bonferroni correction when investigating a single context, and may use FDR within a context when investigating multiple contexts [4, 15]. After conducting an association test between a phenotype and genetically regulated gene expression, an additional Bonferroni correction is often employed across all tested expression-context-phenotype trios [15]. As this approach across all expression-context-phenotype trios may be too stringent, FDR may also be used. However, adjusting for the FDR within each context or across all contexts simultaneously may lead to an inflation or deflation to the false discovery proportion within certain contexts [17].

To simultaneously control the FDR across all contexts at once, a hierarchical false discovery correction—treeQTL—was developed [17]. Though treeQTL was originally developed for use in eQTL studies, its properties hold for any false discovery correction where such a hierarchy (e.g. gene level and gene-context level) exists[18]. Briefly, TreeQTL first combines all gene-context p-values for a given gene simultaneously using Simes’s procedure (other related procedures may also be used) to determine if there is an association at this given locus. If there is an association at the locus, FDR is then employed across the contexts within that gene. Importantly, if a gene does not have a significant association as determined by the first step, contexts are not included in the additional correction procedure, thus decreasing the number of tests that need to be accounted for in multiple correction. This approach has been shown to properly control the false discovery rate across an arbitrary number of contexts and levels in the hierarchy, making it an invaluable tool in the context of gene, gene-context, and gene-context-trait discoveries.

To properly adjust the FDR for CONTENT, we use a hierarchy of 3 levels; (1) at the level of the gene, (2) at the level of the context, and (3) at the level of the method or model.

**Figure S3.**
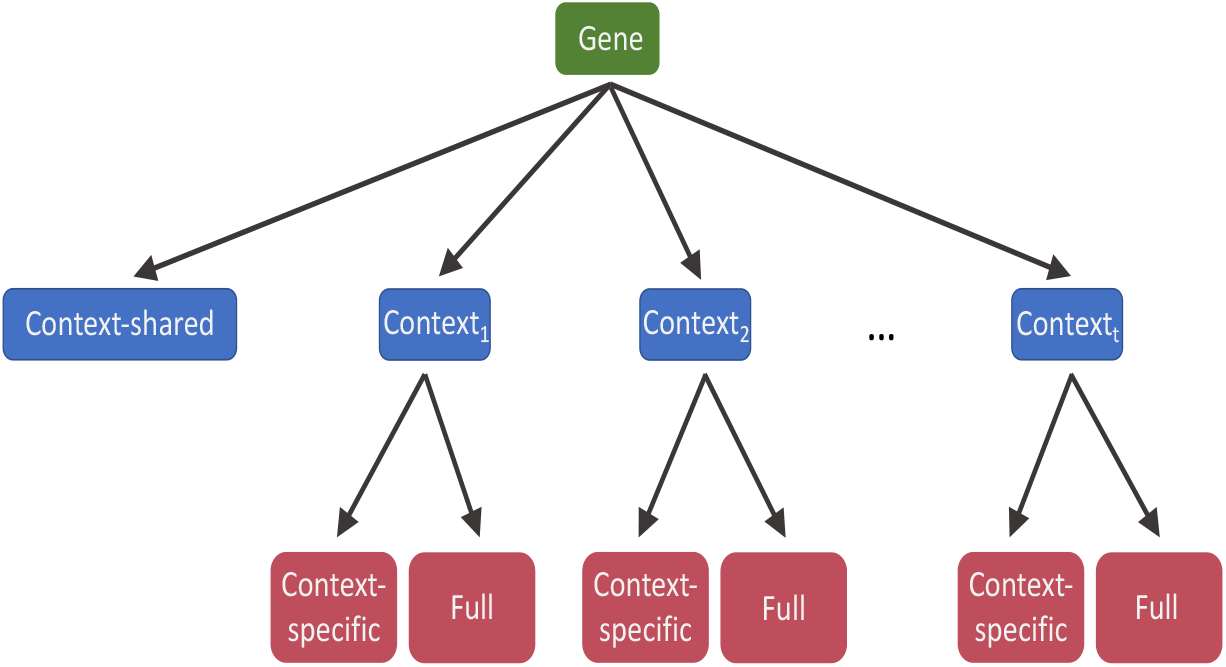
Hierarchical false discovery correction. Here, we show the structure of the hypothesis tests for determining whether a gene has a heritable component. A gene (green, top level) is considered heritable if it has a heritable context-shared component or if it was heritable for a specific context (blue, second level). A given gene-context may be heritable due to either the full or context-specific model of CONTENT (red, third level).

**Figure S4.**
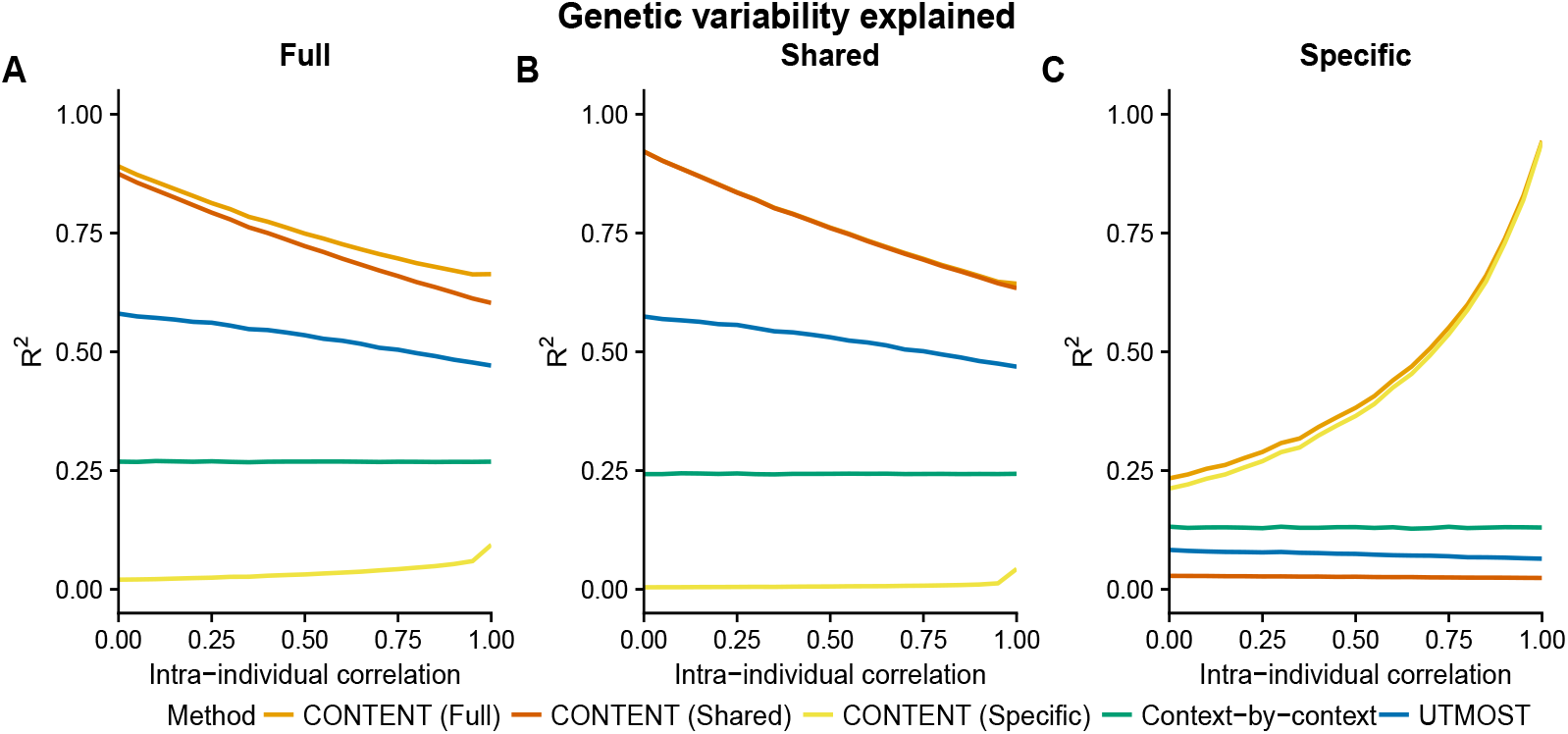
CONTENT is powerful and well-calibrated in simulated data. Accuracy of each method to predict the genetically regulated gene expression of each gene-context pair for different correlations of intra-individual noise across contexts. Mean adjusted *R*^2^ across contexts between the true (A) full, (B) shared, and (C) specific genetic components of expression and the predicted component for each method and for different levels of intra individual correlation. We show here the accuracy for each component and method for all gene-contexts pairs, regardless of whether they had only context-shared or had both context-shared and context-specific effects. Notably, 75% of gene-contexts did not have a context-specific effect, and therefore CONTENT(Shared) captures nearly all of the full variability in these contexts (i.e. the full model is comprised of only shared effects). Further, as only 25% of gene-contexts had context-specific effects, CONTENT(Specific) on average captures very little of the full variability.

### Simulations under additional parameter settings

In this section, we evaluate CONTENT, UTMOST, and the context-by-context approach using the same simulations framework as in the main text (Figure 2), however here we show each methods’ performance while varying additional parameters (Figure S5). We also show the performance of each method when the heritability of the context-shared and context-specific effects are equal (.2; Figure S6) and where the context-shared heritability is less than the context-specific effects (.1 and .3 respectively; Figure S7)).

**Figure S5.**
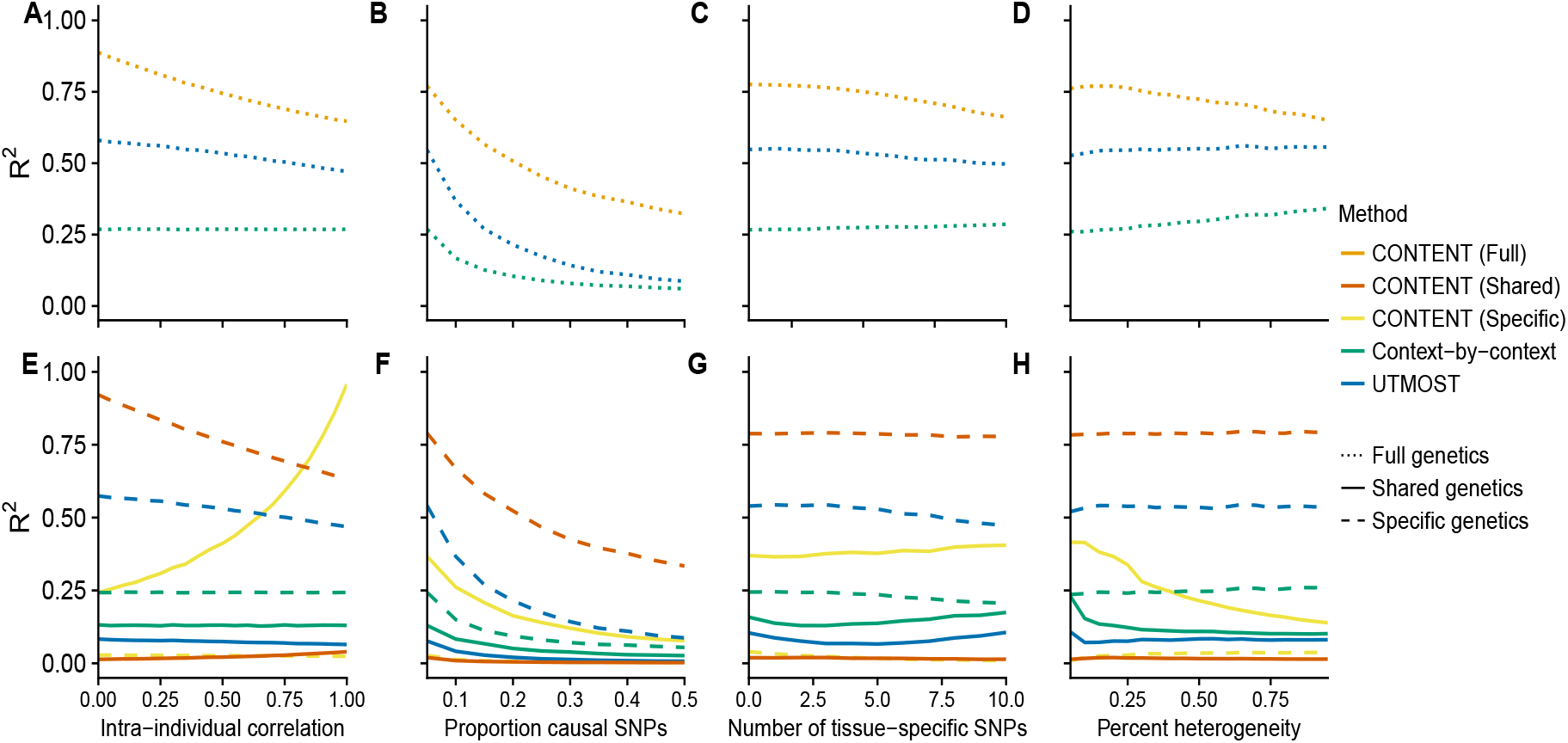
Prediction accuracy across simulated data with higher context-shared than context-specific heritability (.3 and .1 respectively). Under a simulations framework, we evaluated the performance of each method to predict the total expression using the mean adjusted *R*^2^ for each gene-context pair across all iterations for different (A,E) correlation between contexts, (B,F) proportion of causal cis-SNPs, (C,G) number of context-specific SNPs, and (D,H) the percent of contexts with context-specific effects on top of the shared effects. (A-D) show the correlation between the true full (specific + shared) genetic component and the estimated full genetic component of each method, and (E-H) show the correlations of the true genetic shared and specific genetic components of the output of each method (where CONTENT separates the two).

For all methods, the baseline of parameters was .3 shared heritability, .1 specific heritability, 500 cis-SNPs, 20 contexts, 0 correlation between contexts, .05 percent causal SNPs, 2 context-specific SNPs, and 20% specificity (signifying the overlap with the shared effects, as well as the percent of contexts with a specific effect). CONTENT continued to outperform the previous methods, and UTMOST consistently outperformed the context-by-context approach. UTMOST consistently performed better than the context-by-context approach, likely as this simulation framework better fits the model’s assumptions. We note that UTMOST performed better than CONTENT when there were context-specific effects across all contexts (and this set of effects lied on top of SNPs with a shared effect) and the heritability of context-specific effects dominated the heritability of context-shared effects (Figure S7). Given our analysis of GTEx data this architecture may not be entirely common, however this provides further evidence that each method may outperform the other under different architectures, and should therefore be used in complement with the others.

**Figure S6.**
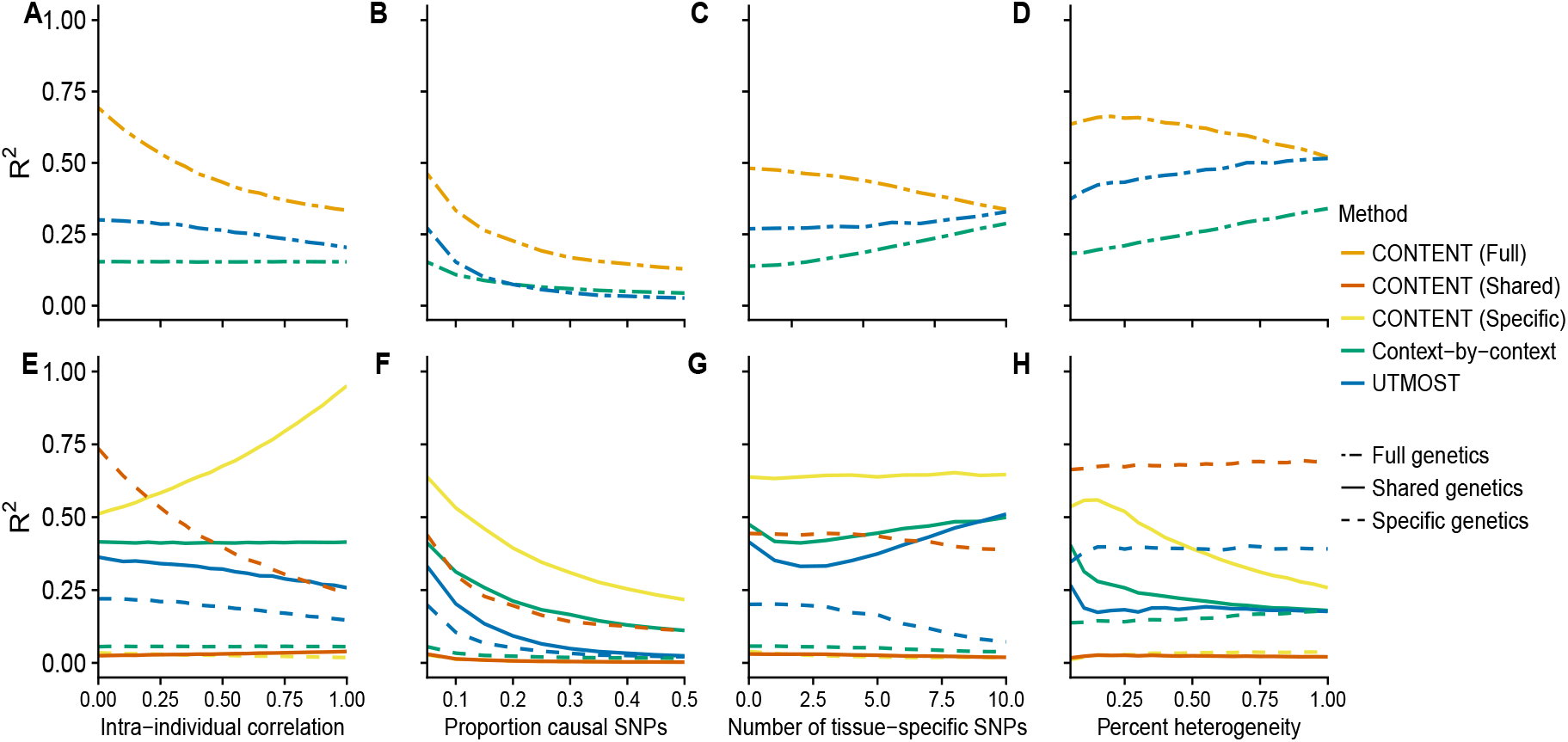
Prediction accuracy across simulated data with equal context-shared and context-specific heritability (.2). Under a simulations framework, we evaluated the performance of each method to predict the total expression using the mean adjusted *R*^2^ for each gene-context pair across all iterations for different (A,E) correlation between contexts, (B,F) proportion of causal cis-SNPs, (C,G) number of context-specific SNPs, and (D,H) the percent of contexts with context-specific effects on top of the shared effects. (A-D) show the correlation between the true full (specific + shared) genetic component and the estimated full genetic component of each method, and (E-H) show the correlations of the true genetic shared and specific genetic components of the output of each method (where CONTENT separates the two).

**Figure S7.**
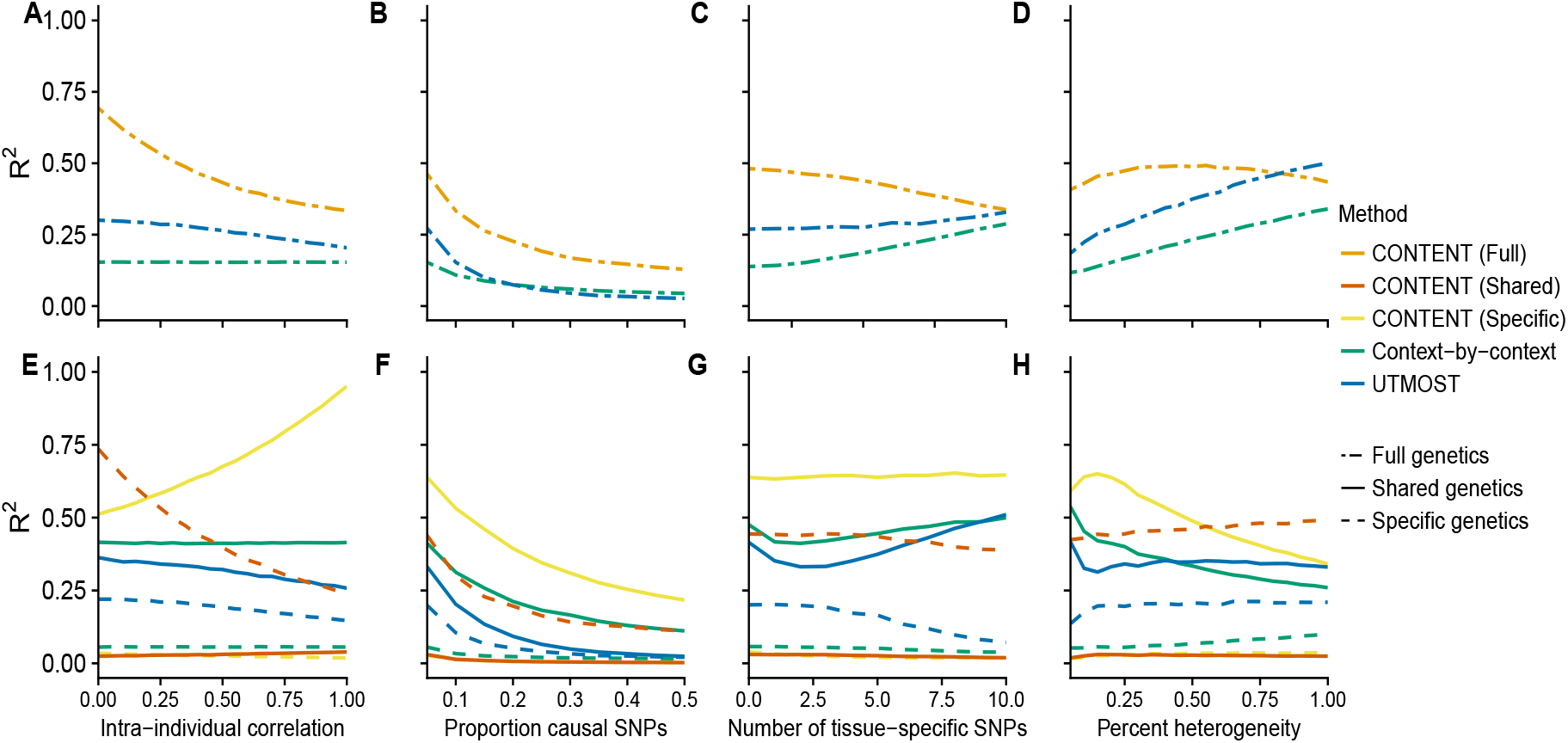
Prediction accuracy across simulated data with lower context-shared than context-specific heritability (.1 and .3 respectively). Under a simulations framework, we evaluated the performance of each method to predict the total expression using the mean adjusted *R*^2^ for each gene-context pair across all iterations for different (A,E) correlation between contexts, (B,F) proportion of causal cis-SNPs, (C,G) number of context-specific SNPs, and (D,H) the percent of contexts with context-specific effects on top of the shared effects. (A-D) show the correlation between the true full (specific + shared) genetic component and the estimated full genetic component of each method, and (E-H) show the correlations of the true genetic shared and specific genetic components of the output of each method (where CONTENT separates the two).

**Figure S8.**
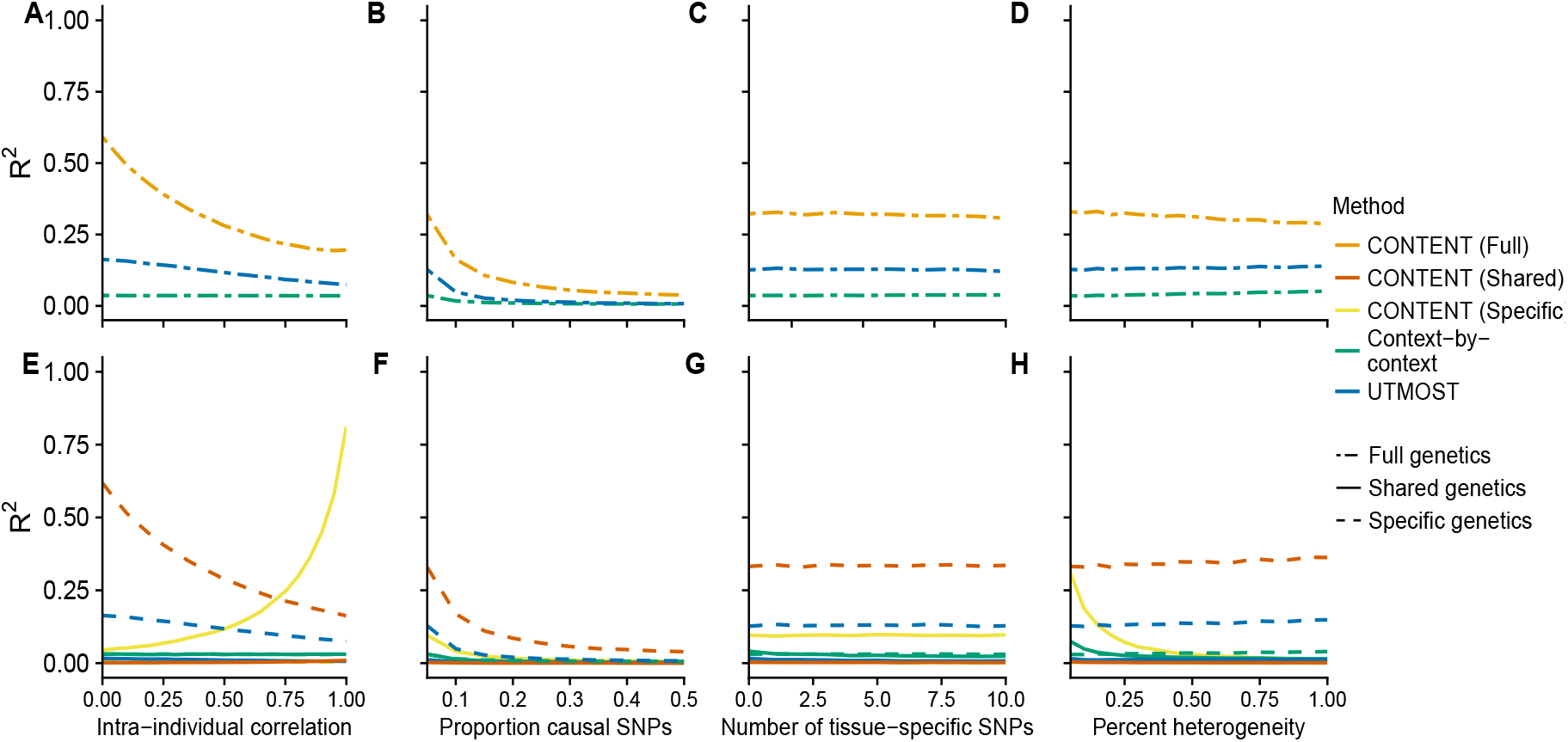
Prediction accuracy across simulated data (2,000 cis-SNPs). Under a simulations framework, we evaluated the performance of each method to predict the total expression using the mean adjusted *R*^2^ for each gene-context pair across all iterations for different (A,E) correlation between contexts, (B,F) proportion of causal cis-SNPs, (C,G) number of context-specific SNPs, and (D,H) the percent of contexts with context-specific effects on top of the shared effects. (A-D) show the correlation between the true full (specific + shared) genetic component and the estimated full genetic component of each method, and (E-H) show the correlations of the true genetic shared and specific genetic components of the output of each method (where CONTENT separates the two).

### Runtimes of methods

We compared the runtimes and memory requirements of our software that fits both CONTENT and the context-by-context approach (10-fold cross-validation) to UTMOST (5-fold cross-validation). Our software takes advantage of the memory-mapped,fast penalized linear regression framework implemented by R package `bigstatsr` [71]. When we tested both approaches on 100 randomly-selected GTEx genes, not only was the runtime of UTMOST—while running half as many cross-validation folds as our method—on average over 3x the runtime of running our software, but the average memory required by UTMOST was also over 10x the memory required by our software.

**Figure S9.**
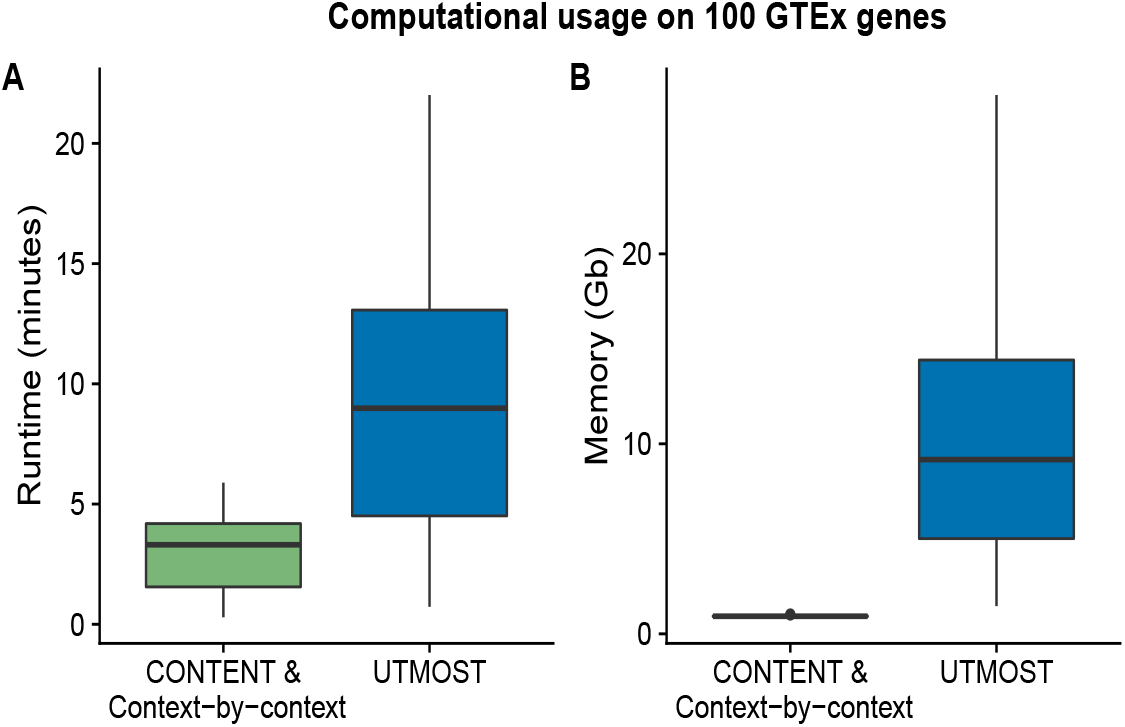
Runtime and memory usage of CONTENT and the context-by-context approach compared to UTMOST. We saved the runtime and memory usage for UTMOST and our software that fits both CONTENT and the context-by-context approach on 100 randomly-selected GTEx genes. The average runtime and memory usage of running UTMOST was over 3x and 10x the runtime and memory usage of running our software that fits both CONTENT and the context-by-context approach.

**Figure S10.**
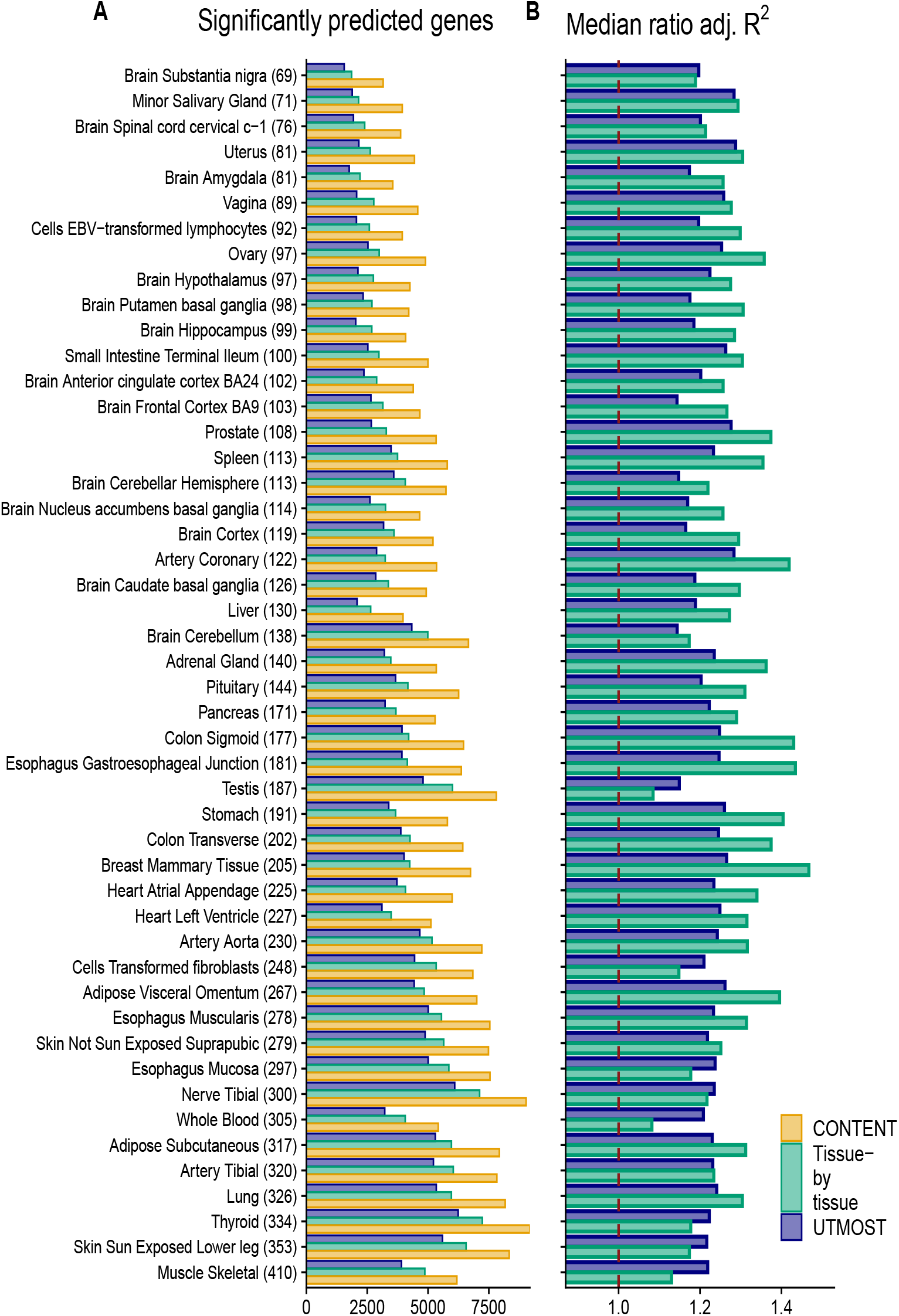
Power of CONTENT, UTMOST and the context-by-context model across GTEx on genes run by UTMOST. (A) The number genes of genes with a significantly predictable component across each context with sample size included in parentheses (B) The median ratio of adjusted *R*^2^ (CONTENT/context-by-context,CONTENT/UTMOST) across the union of genes significantly predicted by CONTENT and either the context-by-context model or UTMOST.

**Figure S11.**
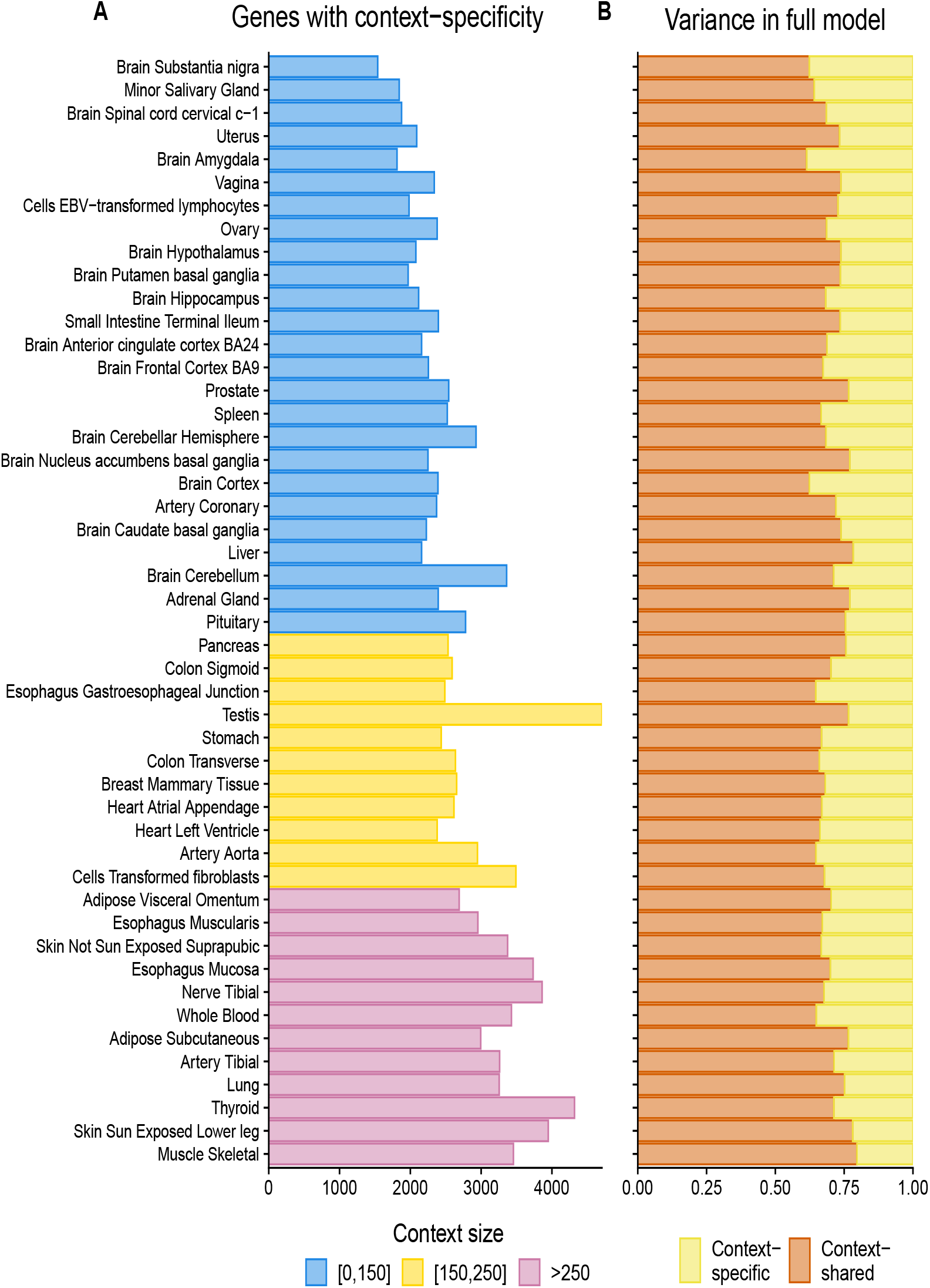
PContribution of context-specific genetic regulation in GTEx. (A) The number of genes with a significant (FDR ≤ 5%) CONTENT(Specific) model of expression in GTEx. (B) Proportion of expression variance of CONTENT(Full) explained by CONTENT(Specific) and CONTENT(Shared) for genes with a significant CONTENT(Full) model.

### Evaluation TWAS simulations and fine-mapping

In this section, we explore the ability of each method to correctly determine the gene-context pair responsible for the association with the phenotype in TWAS. Notably, in these simulations we limited our analyses to situations in which the causal context(s) has been observed. In real data applications, this may not occur, and in such cases, further complexities may arise due to genetic correlation. In these situations, it is likely that all methods will produce false-positive gene-context associations since the true causal context is missing. The complexities posed by missing contexts and cell-types are beyond the scope of this manuscript, and we leave the development of relevant methodology as future work.

Importantly, the models built by CONTENT(Full) can be explained by either the context-shared component, the context-specific component, or both. To implicate a genuine CONTENT(Full) gene-context association (i.e., to elucidate whether a specific context’s expression is more strongly associated than the context-shared expression), we propose using only gene-context pairs whose CONTENT(Full) TWAS test statistic is greater in magnitude than the context-shared TWAS test statistic—termed “CONTENT(Fine).” In our simulations we used a test statistics threshold of .5 and found that this heuristic controlled the false positive rate of the CONTENT(Fine) model’s associations as well as enriched for correctly-associated contexts.

We evaluated the ability of each method to implicate the correct eAssociation in simulated TWAS data. Across a range of heritability and hetereogeneity (percent of contexts with context-specific genetic effects in addition to the main effects), we simulated 1000 genes for 20 contexts, 100 of which had 3 contexts whose genetic component of expression was associated with the phenotype. We considered sensitivity and specificity as the ability of each method to implicate the correct context for an associated gene. To evaluate sensitivity and specificity, we examined which gene-context pairs were significantly associated with the phenotype after employing the hierarchical false discovery correction [17] as the gene-based false positive rate was well-controlled across methods using this approach.

In the absence of context-shared genetic effects, all methods showed high specificity and sensitivity (Figure S13). However, as the genetic variability became more context-shared, the specificity and sensitivity of the context-by-context approach and UTMOST dropped substantially (Figure S13). As neither the context-by-context approach nor UTMOST attempt to deconvolve the context-shared and context-specific effect sizes, their weights for a given context contain both context-shared and context-specific signal. Thus when the context-shared effects dominate the heritability, both methods are likely to suggest context-specific associations across all contexts that express an associated gene. The specificity of CONTENT’s context-specific component, as well as the full model’s weighting of each expression component are paramount to its specificity and sensitivity, as shown by its robust performance across various mixtures of genetic effects (Figure S13).

**Figure S12.**
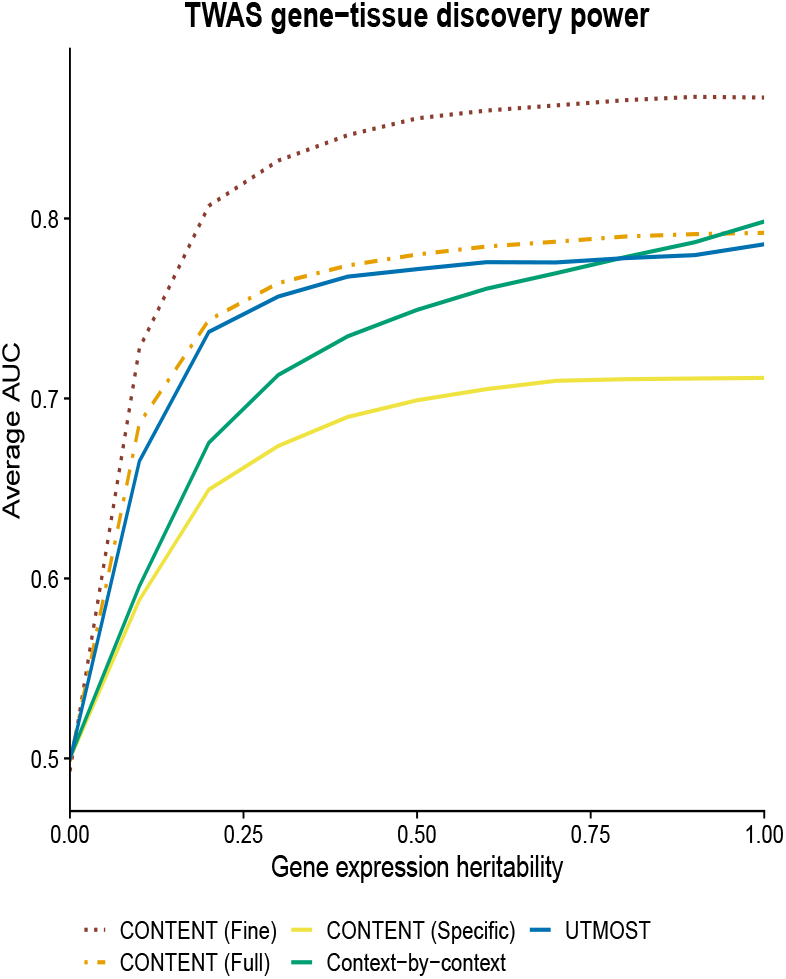
Using a heuristic to fine-map CONTENT(Full) associations. Average AUC from 1000 TWAS simulations while varying the overall heritability of gene expression. Each phenotype (1000 per proportion of heritability) was generated from 300 (100 genes and 3 contexts each) randomly selected gene-context pairs’ genetically regulated gene expression, and the 300 gene-context pairs’ genetically regulated expression accounted for 20% of the variability in the phenotype.

In the GTEx dataset, the fine-mapping TWAS associations produced by our heuristic for the CONTENT(Full) model produced broad associations across many tissues. Though we observed many correct fine-mapping associations for several known gene-trait etiologies (e.g. CYLD and esophagus mucosa in Crohn’s [74], LIPC and liver in HDL [75], SORT1 in liver in LDL and HDL [76–78]), there was not consistent enrichment of a specific tissue known to be relevant for a given trait (for example, the pancreas was not over-represented in associations of Type 2 Diabetes). This could be because the correct tissue or context is missing from the data, horizontal or vertical pleiotropy, or other unknown reasons. As the fine-mapping heuristic performed well in simulated data under a known architecture and where all contexts are observed, we are hopeful that the context-specific estimates will be useful in downstream tissue fine-mapping methods.

**Figure S13.**
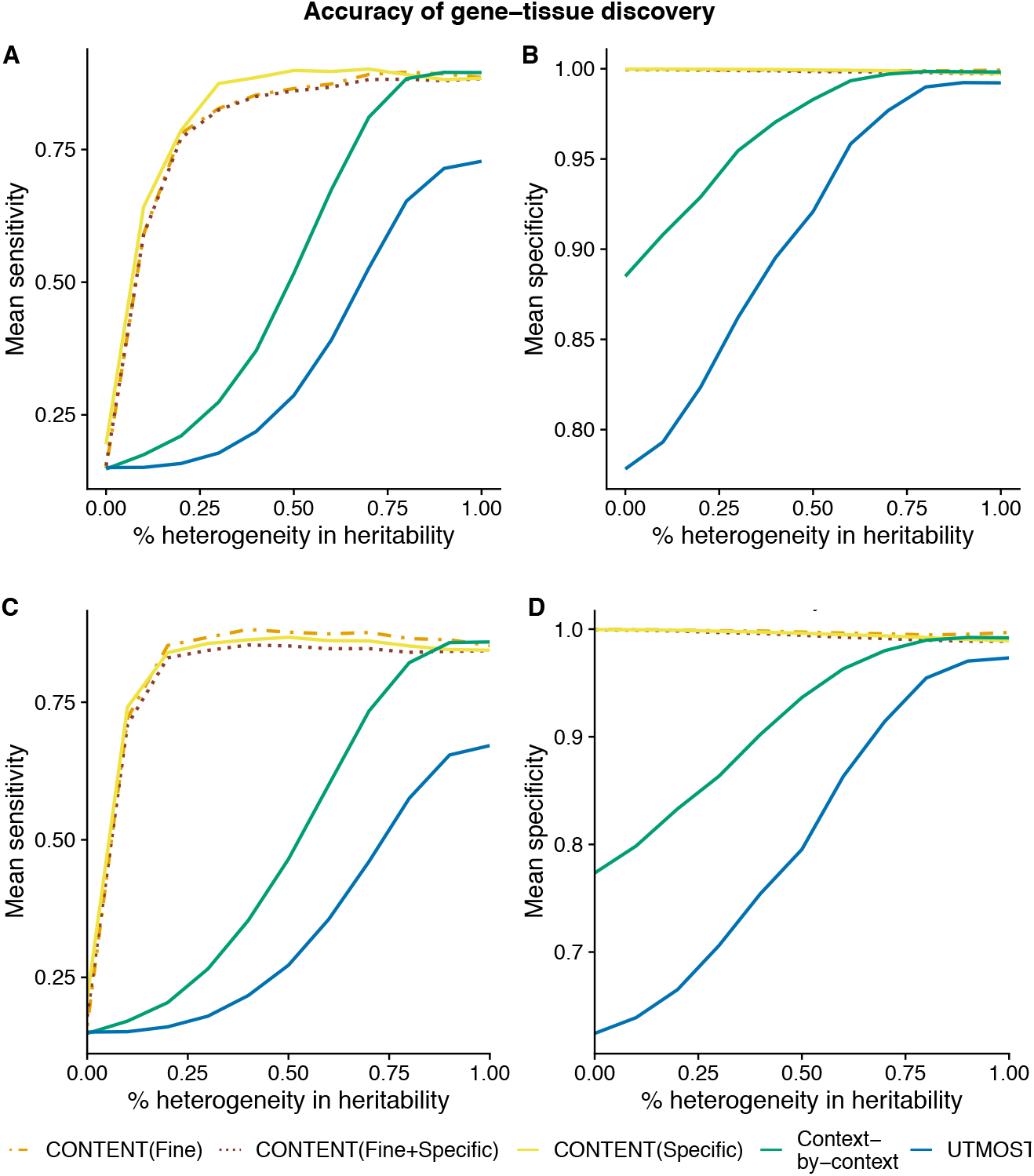
CONTENT is sensitive and specific. We simulated 1000 phenotypes from 300 randomly selected gene-tisue pairs’ expression while varying the percent heterogeneity and performed a TWAS using the weights output by each method. (A,B) When the total proportion of variability in the phenotype due to the genetically regulated gene expression is .5 and (C,D) when the proportion is .2. The full model of CONTENT was the most sensitive when finding the correct gene-context pair, and is most powerful when there is non-negligible context-specific heritability in addition to the tissue-shared heritability.

**Table S1.** GWAS summary statistics used as input for TWAS. Abbreviation used for each trait as well as and its respective study and sample size. The collection of traits from the UKBiobank were self-reported and measured on the same set of individuals across traits.

| Symbol | Trait | Study | Sample Size |
| --- | --- | --- | --- |
| AD | Alzheimer's disease | Lambert et al. Nat Genet. 2013 | 74,046 |
| Asthma | Asthma (self-reported) | UKBB Loh et al. 2018 Nat Genet | 361,141 |
| Bipolar | Bipolar Disorder | PGC Cell 2018 | 73,684 |
| CAD | Coronary Artery Disease | CARDIoGRAM Nat Genet. 2011 | 86,995 |
| CKD | Chronic Kidney Disease | Wuttke et al. Nat Genet. 2019 | 1,046,070 |
| Crohn's | Crohn's Disease | IIBDGC Europeans Nat Genet. 2015 | 13,974 |
| Eczema | Eczema (self-reported) | UKBB Loh et al. 2018 Nat Genet | 361,141 |
| FastGlu | Fasting Glucose | MAGIC Nat Genet. 2012 | 96,496 |
| HDL | High-density Lipoprotein | Teslovich et al. Nature 2010 | 99,900 |
| IBS | Irritable bowel syndrome (self-reported) | UKBB Loh et al. 2018 Nat Genet | 361,141 |
| LDL | Low-density lipoprotein | Global lipids genetics consotrium Nat Genet 2013 | 188,577 |
| Lupus | Systemic Lupus Erythromous | Bentham et al. Nat Genet 2015 | 23,210 |
| MDD | Major Depression Disorder | PGC; Howard et al. Nat Neuro 2019 | 807,553 |
| MS | Multiple Sclerosis (self-reported) | UKBB Loh et al. 2018 Nat Genet | 361,141 |
| PBC | Primary biliary cirrhosis | Cordell et all. Nat Comm 2015 | 13,239 |
| Psoriasis | Psoriasis (self-reported) | UKBB Loh et al. 2018 Nat Genet | 361,141 |
| RA | Rheumatoid Arthritis | Okada et al. Nature 2013 | 103,638 |
| Sarcoidosis | Sarcoidosis (self-reported) | UKBB Loh et al. 2018 Nat Genet | 361,141 |
| Sjogren | Sjogren's Syndrome (self-reported) | UKBB Loh et al. 2018 Nat Genet | 361,141 |
| T1D | Type 1 Diabetes | Inshaw et al. Diabetologia 2021 | 17,685 |
| T2D | Type 2 Diabetes | DIAGRAM Nat Genet 2018 | 898,130 |
| Ulc colitis | Ulcerative Colitis (self-reported) | UKBB Loh et al. 2018 Nat Genet | 361,141 |

### TWAS discoveries as a function of heritability thresholding

In the main text, we put forth all gene-context pairs that were genetically predicted with a nominal pvalue of .1. As the procedure we use for false discovery adjustment was robust across contexts, we evaluated the number of discoveries that are potentially made when raising the threshold for the nominal pvalue. Our results suggest that there may be minimal correlation between genetic-predictability and strength of TWAS association.

**Figure S14.**
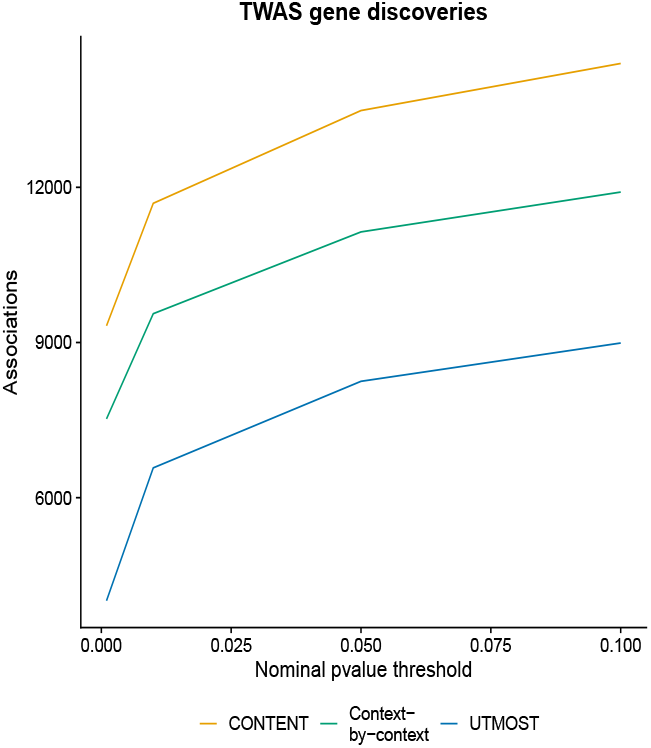
TWAS discoveries across predictability thresholds. The number of hierarchical-FDR-corrected TWAS discoveries as a function of the nominal pvalue cutoff for a given gene-tissue’s cross-validation expression prediction.

### CONTENT can accommodate additional levels of pleiotropy among contexts

While the original model of CONTENT enables a simple decomposition into a component that is shared across all contexts and another that is specific to a single context, there may be cases in which additional sharing exists across a subset of contexts. For example, the group of brain tissues measured in the GTEx consortium have shown similar patterns in terms of cis-genetic variability [2, 25, 79] as well as intra-individual residual correlations (Figure S1). To further disentangle the shared and tissue-specific genetic components of expression in the brain tissues, we added an additional term to the CONTENT decomposition which accounts for genetic effects that are only shared across the brain tissues. In more detail, we decompose the original context-shared component of expression into a new context-shared component that is shared across all tissues and a brain-shared component that is shared across only the brain tissues:

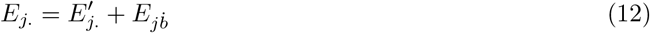

Here, 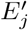. (the new context-shared term) is an intercept, 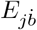 (the brain-shared term) is the effect size on an indicator variable for brain tissues, and estimates of both terms are generated for each individual using a simple linear regression. While introducing an additional term for the shared component will increase the resolution of the model, i.e. the novel model may discover new components of brain-sharing that were miscategorized as tissue-specific in multiple brain tissues, there may be a significant loss in power as this decomposition is only possible for individuals who have been sampled in both multiple brain and non-brain tissues. Additionally, under this decomposition, the full model for brain tissues contains three terms—the context-specific, brain-shared, and globally shared—resulting in a loss of a degree of freedom relative to the original model.

To evaluate the effect of an additional source of effects-sharing on the performance of CONTENT, we simulated an additional genetic effect that lied on top of a subset of SNPs with a main, overall context-sharing effect in 25% of the contexts. As the heritability of this additional source of sharing grew, the context-specific component of CONTENT began to capture variability due to both the context-specific and secondary context-shared effects (Figure S15). When we used CONTENT brain, the context-specific component of CONTENT no longer produced predictors that captured variability due to the additional source of effects-sharing (mean *R*^2^ of true brain effects and predicted tissue-specific effects dropped from 0.127 to 0.004 across simulations), and the component responsible for capturing the additional source of effects-sharing–CONTENT(Brain)– was robust (average *R*^2^ between true and predicted brain-shared effects 0.49).

We applied the CONTENT brain model to GTEx, but note that such a component is only identifiable for individuals who have been sampled in both multiple brain and multiple non-brain tissues. For our analysis of the GTEx data, our sample size decreased to 12,904 genes, 26 tissues, and 150 individuals when using CONTENT brain. In general, using this model, the number of genetic tissue-specific components in the brain tissues decreased (Figure S16). Of the genes that were implicated in the original CONTENT model as having a tissue-specific component but were no longer captured in the CONTENT brain model with a tissue-specific component, roughly 12% overlapped with the genes implicated by the additional brain-shared component. The CONTENT brain model discovered 4,811 genes with an overall tissue-shared component as well as 1,960 genes with a brain-shared components (of which 66% also had an overall tissue-shared component). The prediction accuracy was similar in both the original and brain models of CONTENT (Figure S17).

**Figure S15.**
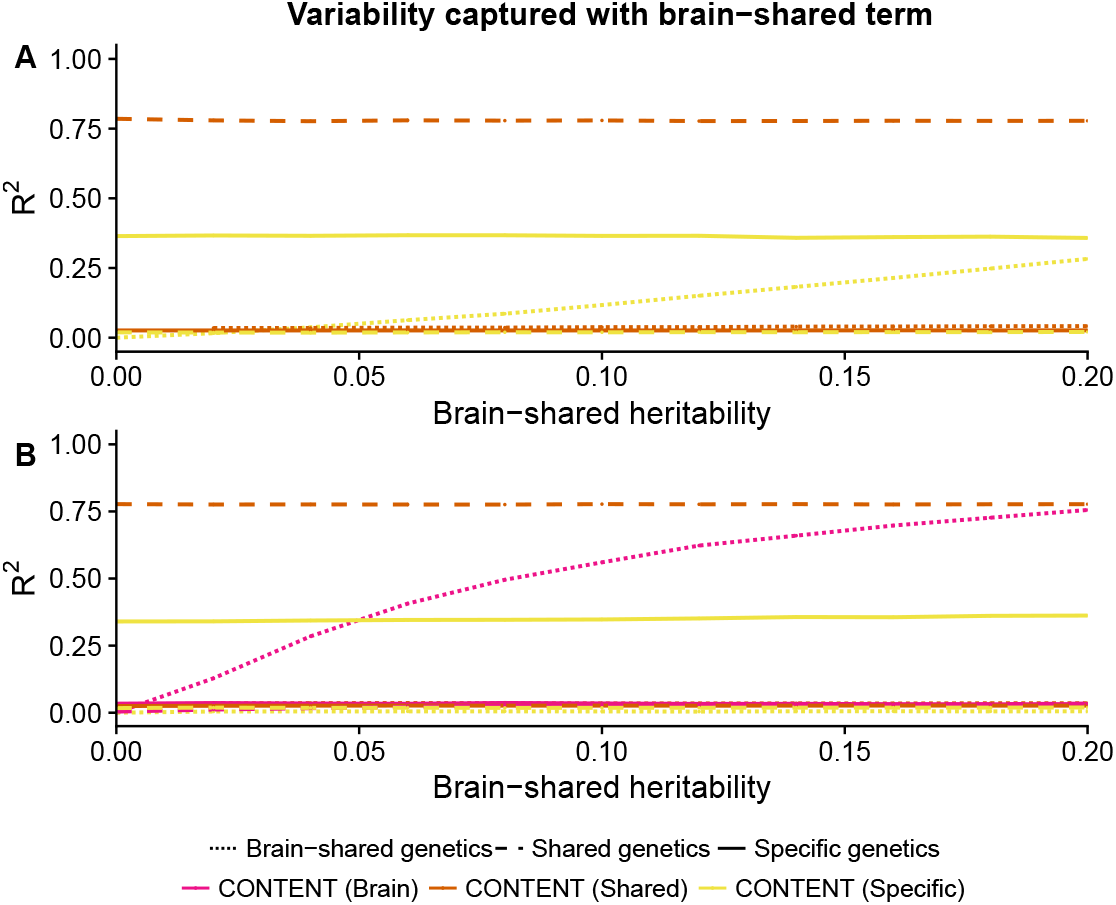
Additional sources of tissue-sharing may confound the tissue-specific component. (A) The original CONTENT model without accounting for the additional source of shared genetic effects when such a component exists. (B) When we introduce an additional shared component to the CONTENT model, CONTENT(Brain), the specific component does not capture this additional component, and the additional component is recovered.

We next compared the performance of the original CONTENT model to the CONTENT brain model in TWAS using simulated data (generated as aforementioned) as well as GTEx. While the mean AUC between both methods was similar in the simulated data, CONTENT brain was more sensitive than the original CONTENT model when shared brain effects existed (Figure S18). Further, despite the fact that the sample size and number of tissues in GTEx data subsetted for the brain model is smaller, CONTENT discovered a non-trivial additional number of TWAS associations (Figure S19). In several neurological disorders, the number of context-specific genes decreased when using the brain model, however the brain model discovered genes whose genetics were shared across only the brain-shared component (Figure S19). When we examined previous TWAS associations, such as APOC1 and AD, the original CONTENT approach showed association with the thyroid. However, this signal was removed using the brain-pleiotropy approach and the brain pleiotropic component showed significant association (p=2.20e-23). We observed a similar trend with APOE, where the original CONTENT model implicated several brain tissue associations but no significant shared association. The brain pleiotropy model in turn discovered a brain-tissue-shared component with significant evidence of association (p=2.47e-29). Both genes are known to have neuronal roles in Alzheimer’s disease [80].

### Performance in GTEx when using the brain component

We ran the original and brain versions of the CONTENT model on 12904 genes in 26 tissues and 150 individuals in the GTEx dataset. These individuals were measured in at least 3 brain and non-brain tissues. Interestingly, each model discovered eGenes that were not discovered by their counterpart. The amount of variability was roughly the same in both versions of the model, but the adjusted *R*^2^ was slightly higher in non-brain tissues and slightly lower in brain tissues in the brain model. Importantly, the brain tissues in the brain model have 3 explanatory variables and therefore suffer a larger penalty in the adjusted *R*^2^ relative to the original CONTENT model. The adjusted *R*^2^ improved in the non-brain tissues however, suggesting that the context-shared and context-specific components may be less confounded by the brain tissues in the brain model than in the original model.

**Figure S16.**
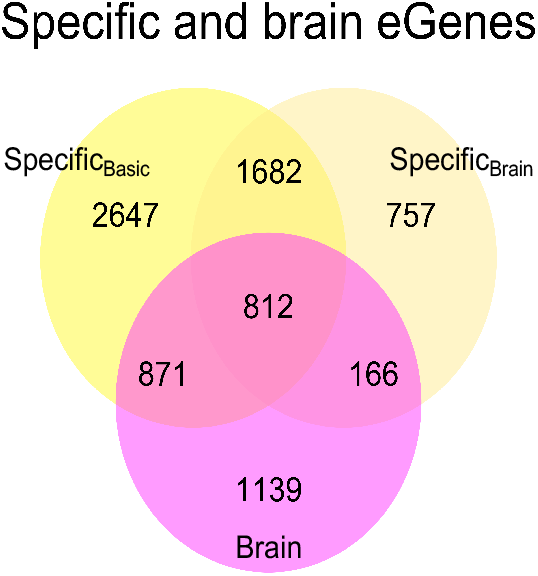
Additionally sources of effects-sharing may confound the context-specific component. When we run the original CONTENT model and the CONTENT model with the brain-sharing on GTEx genes that are expressed in at least 3 brain and 3 non-brain tissues, many of the previous genetic context-specific components in the brain tissues are absorbed by the additional brain-sharing across brain tissues.

**Figure S17.**
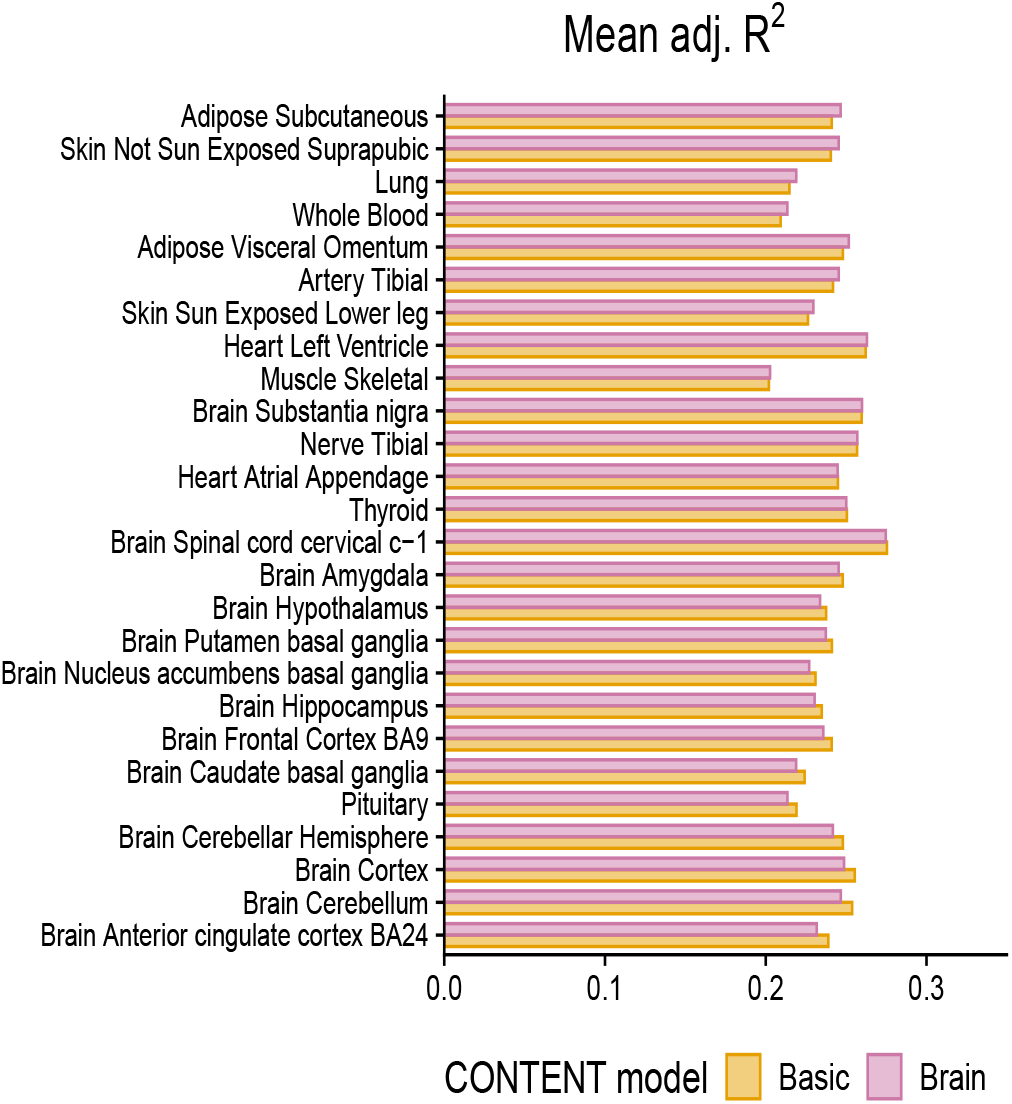
Prediction accuracy across tissues in the brain and original CONTENT model. The difference in adjusted *R*^2^ in the brain and original CONTENT(Full) models. While the variability explained is markedly similar in both versions of the model, the adjusted *R*^2^ generally increased in non-brain tissues, and decreased in the brain tissues in the brain model.

**Figure S18.**
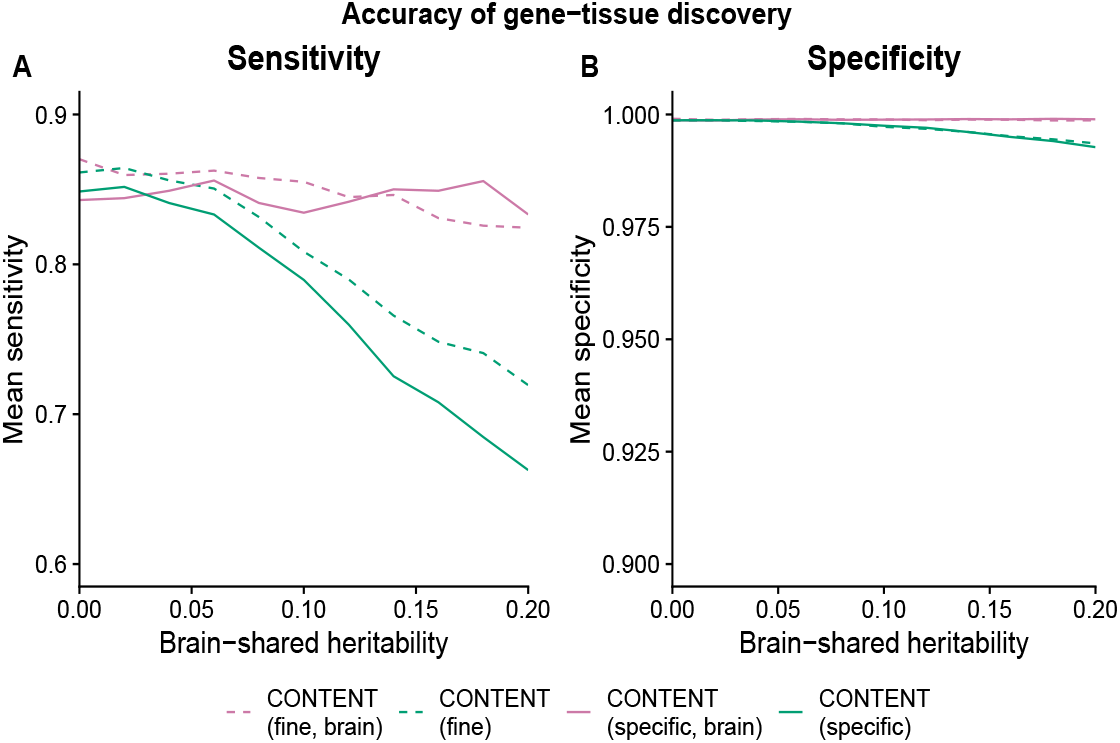
Simulated TWAS with brain-shared genetic effects. While the AUC and specificity of the original CONTENT model (green) and the CONTENT model that accounts for brain-shared effects (pink) were nearly the same, the sensitivity was improved when using the brain version of CONTENT in simulated TWAS where there exists brain-shared effects.

### TWAS eGenes discovered using the brain version of CONTENT

We performed TWAS using weights trained by the original and brain versions of the CONTENT model on 26 tissues, 12,094 genes, and 150 individuals in the GTEx dataset for 17. These individuals were measured in at least 3 brain and non-brain tissues, leading the sample size to be smaller than when using the total GTEx data without any such constraint. While the brain version of the CONTENT model discovered more TWAS eGenes than the original model, the brain model discovered fewer context-specific eGenes than the original model.

**Figure S19.**
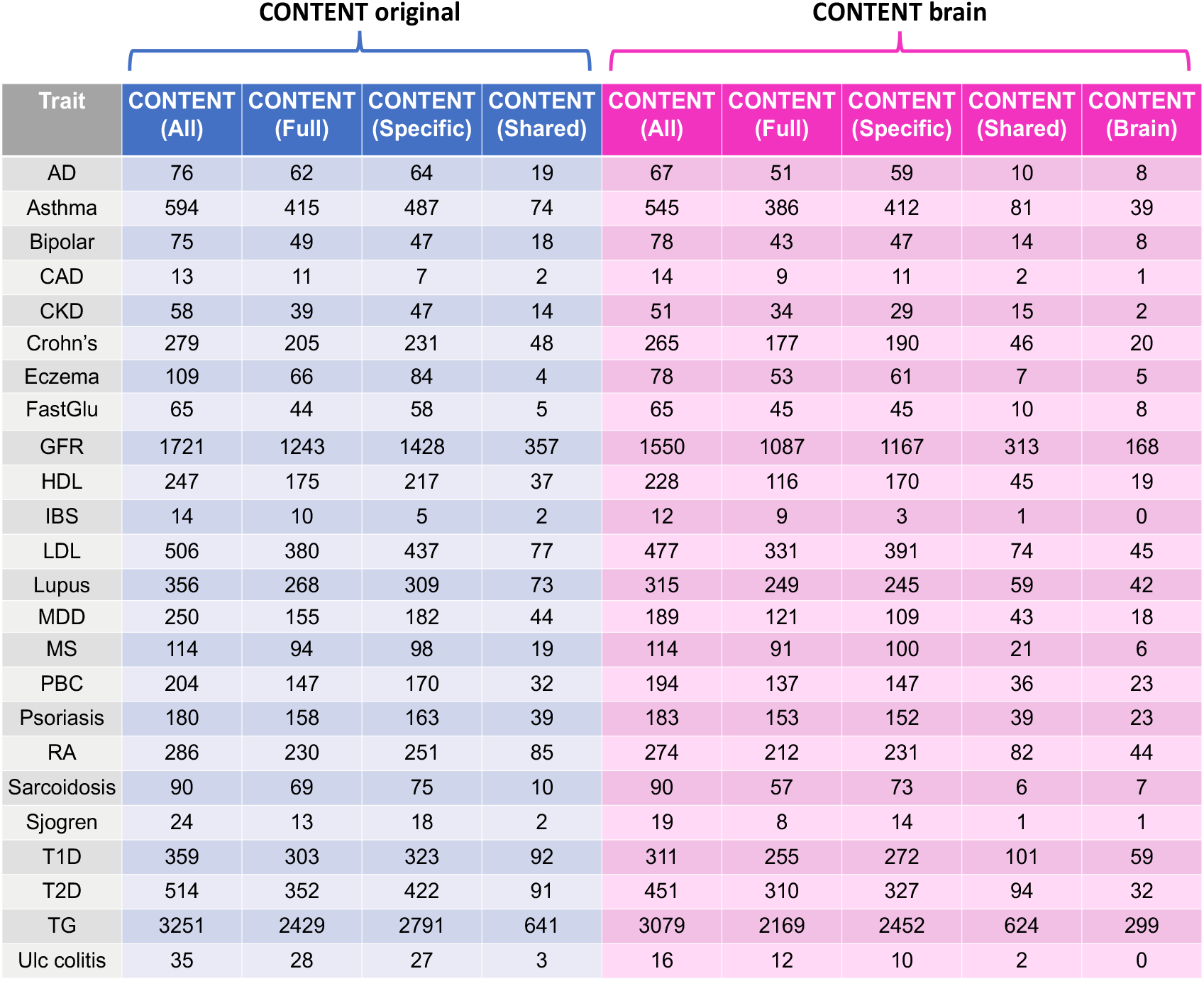
eGenes discovered by each component of CONTENT model in the brain and original models. In total, there were fewer genes discovered using the brain model of CONTENT, however our simulations show that the brain model of CONTENT may improve the resolution of associations.

